# Apical-driven cell sorting optimised for tissue geometry ensures robust patterning

**DOI:** 10.1101/2023.05.16.540918

**Authors:** Prachiti Moghe, Roman Belousov, Takafumi Ichikawa, Chizuru Iwatani, Tomoyuki Tsukiyama, François Graner, Anna Erzberger, Takashi Hiiragi

## Abstract

Tissue patterning coordinates morphogenesis, cell dynamics and fate specification. Understanding how these processes are coupled to achieve precision despite their inherent variability remains a challenge. Here, we investigate how salt-and-pepper epiblast and primitive endoderm (PrE) cells sort and robustly pattern the inner cell mass (ICM) of mammalian blastocysts. Quantifying cellular dynamics and mechanics together with simulations show a key role for the autonomously acquired apical polarity of mouse PrE cells in coupling cell fate and dynamics in tissue contexts. Specifically, apical polarity forms actin protrusions and is required for Rac1-dependent migration towards the ICM surface, where PrE cells are trapped due to decreased tension at their apical domain, while depositing an extracellular matrix gradient, breaking the tissue-level symmetry and collectively guiding their own migration. Tissue size perturbations and comparison with monkey blastocysts further demonstrate that the fixed proportion of PrE/epiblast cells is optimal and robust to variability in embryo size.

## INTRODUCTION

Tissue patterning in developing embryos depends on the coordination between cellular dynamics and fate specification in tissue contexts. While specification of cell fate is typically accomplished through gene regulatory networks activated by secreted morphogens and signalling, dynamic cellular behaviours including division, migration and sorting drive morphogenesis and form spatially organised tissues. In developing tissues, equipotent cells start differentiating to form a heterogeneous mixture of cell populations. Through cell rearrangements and sorting, tissues refine patterns and generate sharp boundaries, such as in the neural tube (Xiong et al. 2013), rhombomeres (Kesavan et al. 2020), somites (Miao et al. 2023; Watanabe et al. 2009), and glandular epithelia (Cerchiari et al. 2015). In these scenarios, local cell rearrangement and cell fate changes can achieve pattern precision. However, how fate specification and dynamics of cells and their inherent variability adapt to the growing, thus changing, tissue size and geometry in order to achieve robust patterning remains less understood. For example, size of the early mouse embryos varies up to 4-fold among *in utero* embryos (Ichikawa et al. 2022) and experimentally manipulated double or half-size embryos develop to term (Tarkowski 1959, 1961), though how they achieve pattern precision remains unclear. While in many organisms, morphogen or signalling gradients extend across tissues and determine the orientation and length-scale of tissue patterns, early mammalian embryos lack such gradients or other forms of pre-patterning (Guo et al. 2010; Rossant and Tam 2009; Yamanaka et al. 2006).

Morphogenesis and patterning of the mouse embryo starts with the formation of a blastocyst, which comprises three distinct cell lineages and a fluid-filled cavity. The trophectoderm (TE) forms the outermost layer of epithelial cells enclosing the pluripotent inner cell mass (ICM) (Yamanaka et al. 2006). ICM cells, initially equivalent and expressing lineage marker genes in a highly heterogeneous manner, progressively differentiate into the innermost embryonic epiblast (EPI) and the cavity-facing, extra-embryonic primitive endoderm (PrE) with the characteristic salt-and-pepper distribution of cell fates (Chazaud et al. 2006; Guo et al. 2010; Ohnishi et al. 2014; Plusa et al. 2008). These cell fates are specified by a gene regulatory network involving lineage-specific transcription factors NANOG and GATA6 and fibroblast growth factor (FGF) signalling (Bessonnard et al. 2014; Frankenberg et al. 2011; Kang et al. 2013; Krawchuk et al. 2013; Schrode et al. 2014). As multiple fluid-filled cavities emerge, expand, coalesce and collapse in the blastocyst, cell sorting within the heterogeneous ICM segregates PrE cells to a monolayer at the cavity surface enveloping the epiblast (Chan et al. 2019; Dumortier et al. 2019; Meilhac et al. 2009; Plusa et al. 2008; Ryan et al. 2019; Saiz et al. 2013, 2020) which is followed by its maturation into a polarised epithelium to pattern the ICM (Gerbe et al. 2008; Saiz et al. 2013). Cellular dynamic mechanisms such as directional movement, cell surface fluctuations, oriented divisions, apoptosis, and positional induction were proposed to drive EPI/PrE segregation (Meilhac et al. 2009; Plusa et al. 2008; Wigger et al. 2017; Xenopoulos et al. 2015; Yanagida et al. 2022), although how these properties arise among ICM cells, the coupling among these processes, and their coordination with cell fate specification and blastocyst morphogenesis are poorly understood.

Thus far, cell fate specification and spatial segregation have been studied independently, and an integrative view of ICM patterning is lacking. Specifically, it remains unclear whether EPI and PrE cells exhibit distinct dynamic movements in the ICM, and if so, what drives them, and how robust patterning is ensured in mouse embryos exhibiting spatio-temporal developmental variabilities, resulting in inevitable variability in embryo size and geometry (Dietrich and Hiiragi 2007; Fabrèges et al. 2023; Ohnishi et al. 2014; Plusa et al. 2008). This is largely due to the technical challenge of analysing cellular dynamics in the presence of the rapidly changing geometry due to the expanding and collapsing fluid cavity. In this study, we systematically analyse cellular dynamics, position, fate, and polarity using reduced systems and blastocyst manipulation to gain mechanistic insights into robust ICM patterning during mouse pre implantation development.

## RESULTS

### Distinct cell movement between EPI and PrE underlies segregation within the ICM

Since the expansion and collapse of the blastocyst cavity rapidly change embryo shape, and make it challenging to track and analyse cellular dynamics, we isolated ICMs from whole blastocysts via immunosurgery (Solter and Knowles 1975) (Figure 1A). This experimental system not only eliminates the abrupt change in overall embryo shape, but also effectively reduces the complexity of cellular dynamics in a three dimensional (3D) and heterogenous geometry to a system near spherical symmetry in which dynamics can be analysed in one radial dimension. In agreement with previous studies (Wigger et al. 2017; Yanagida et al. 2022), we verified that the *in vitro* culture of isolated ICMs faithfully recapitulates the EPI/PrE sorting in the blastocyst in terms of cell number and timing (Figure 1A, B). The total cell number in the ICM was unchanged after immunosurgery (Figure S1A), and those in the ICMs isolated at E3.5 and E4.5, or after 24-hour culture from E3.5, were comparable to ICM cell numbers in E3.5 and E4.5 whole blastocysts, respectively (Figure 1B). We live-imaged the isolated ICMs using a fluorescent reporter of PrE fate, *Pdgfrα^H2B-GFP^,* (Hamilton et al. 2003; Plusa et al. 2008) combined with a ubiquitous *H2B-mCherry* reporter (Abe et al. 2011) (Figure 1C, and Video S1), and quantitatively analysed the dynamics of cell sorting, using a custom, semi-automated nuclear detection and tracking pipeline (Fabrèges et al. 2023) (Figure 1D). To quantify ICM segregation, we define the sorting score as the extent of overlap between EPI and PrE spatial domains (Figure S1B), which describes both live and immunostained ICMs (Figure 1E, Figure S1C).

**Figure 1:**
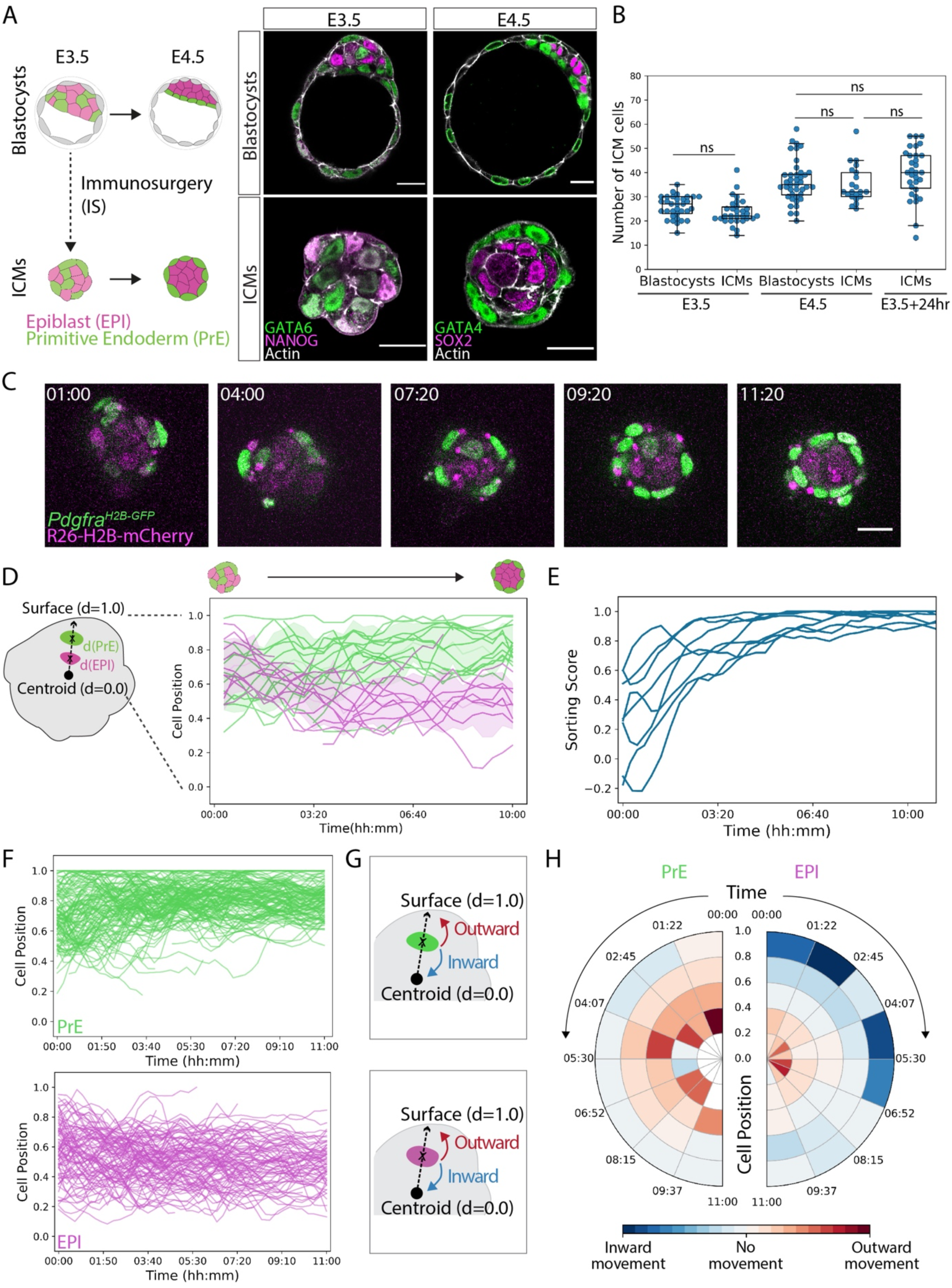
Differential cell movements between epiblast and primitive endoderm contribute to fate segregation in the ICM. A. Schematic representation and immunostaining images of blastocysts and ICMs at stages E3.5 and E4.5. GATA6 and GATA4 are used as markers for PrE fate (green), and NANOG and SOX2 are used as markers for EPI fate (magenta) at stages E3.5 and E4.5, respectively. B. Quantification of total cell numbers in the ICM from blastocysts and isolated ICMs at stage E3.5, blastocysts and isolated ICMs at stage E4.5, and isolated ICMs cultured *in vitro* for 24 hours from stage E3.5 to E4.5. n=33, 30, 40, 21, 31 embryos for the different groups, respectively. Independent samples t-test between E3.5 blastocysts and E3.5 ICMs, *p*=0.106. One-way ANOVA between E4.5 Blastocysts, E4.5 ICMs, and E3.5 ICMs+24hr, *p*=0.145. C. Representative time-lapse imaging of ICMs isolated from E3.5 blastocysts expressing PrE-specific H2B-GFP (*Pdgfrα^H2B-GFP^*, green) and ubiquitous H2B-mCherry (*R26-H2B mCherry*, magenta), out of total 8 datasets from 3 independent experiments. Time is indicated in hh:mm, t=00:00 corresponds to start of live-imaging at stage E3.5+3hours, following completion of immunosurgery. D. Schematic representation of single-cell tracking of EPI and PrE cells from isolated ICMs from (C). Line plots indicating radial distances of all cells from one representative ICM until E4.0 stage. Colour of the line indicates cell fate – PrE, green and EPI, magenta. Shaded regions show spatial dispersion as mean ±SD of cell position along ICM radial axis. The geometric centroid of the ICM is considered as d=0.0 and ICM outer surface is considered as d=1.0 to normalise cell position across samples. Time-series plots for cell position were smoothed using a rolling average. E. Quantification of sorting score for isolated ICMs between stage E3.5 and E4.0. Data from n=8 ICMs. F. Line plots for radial cell position versus time from tracking of PrE and EPI cell movements in isolated ICMs. Time-series plots for cell position were smoothed using a rolling average. Cell tracking data pooled from n=160 PrE cells and n=133 EPI cells from 8 ICMs. G. Schematic diagram for analysis of PrE and EPI cell movements. Cell displacement is measured along the radial axis between consecutive timepoints and classified as inward or outward movement depending on the direction of displacement. H. Polar plots indicating preferential direction of cell movements among PrE and EPI. Cell position is plotted along radial axis, time is plotted along angular axis. Measurements are binned according to initial radial cell position and time. The mean displacement of each interval is plotted, colour indicates direction of movement. Scale bars 20μm. *ns, non-significant*

With these tools established, we analysed the comprehensive cell tracking dataset to examine whether cells exhibit preferential directionality of movement along the ICM radial axis (Figure 1F,G,H). One clearly notable difference was that EPI cells initially on the ICM surface rapidly move inward in early stages of sorting, while surface PrE cells do not show such directed movement (Figure 1G,H). We therefore investigated cellular dynamics at the ICM-fluid interface.

### Apical domain decreases surface tension and positions PrE cells at the fluid interface

To clearly visualise cell shape dynamics, we generated fluorescence-chimeric ICMs by tamoxifen-induced Cre-mediated recombination of the mTmG transgene (Badea, Wang, and Nathans 2003; Muzumdar et al. 2007; Figure 2A). Live-imaging showed that certain cells changed shape and flattened upon reaching the surface (Video S2). To examine if these cells are PrE cells, and if EPI and PrE cells show distinct surface behaviour, we analysed cell shape in immunostained ICMs at E3.5 stage. PrE cells located at the ICM surface were more stretched (Figure 2B), in significant contrast to the more rounded EPI cells (Figure S2A), indicating that it is indeed PrE cells that flatten when reaching the fluid-surface (Figure S2B).

**Figure 2:**
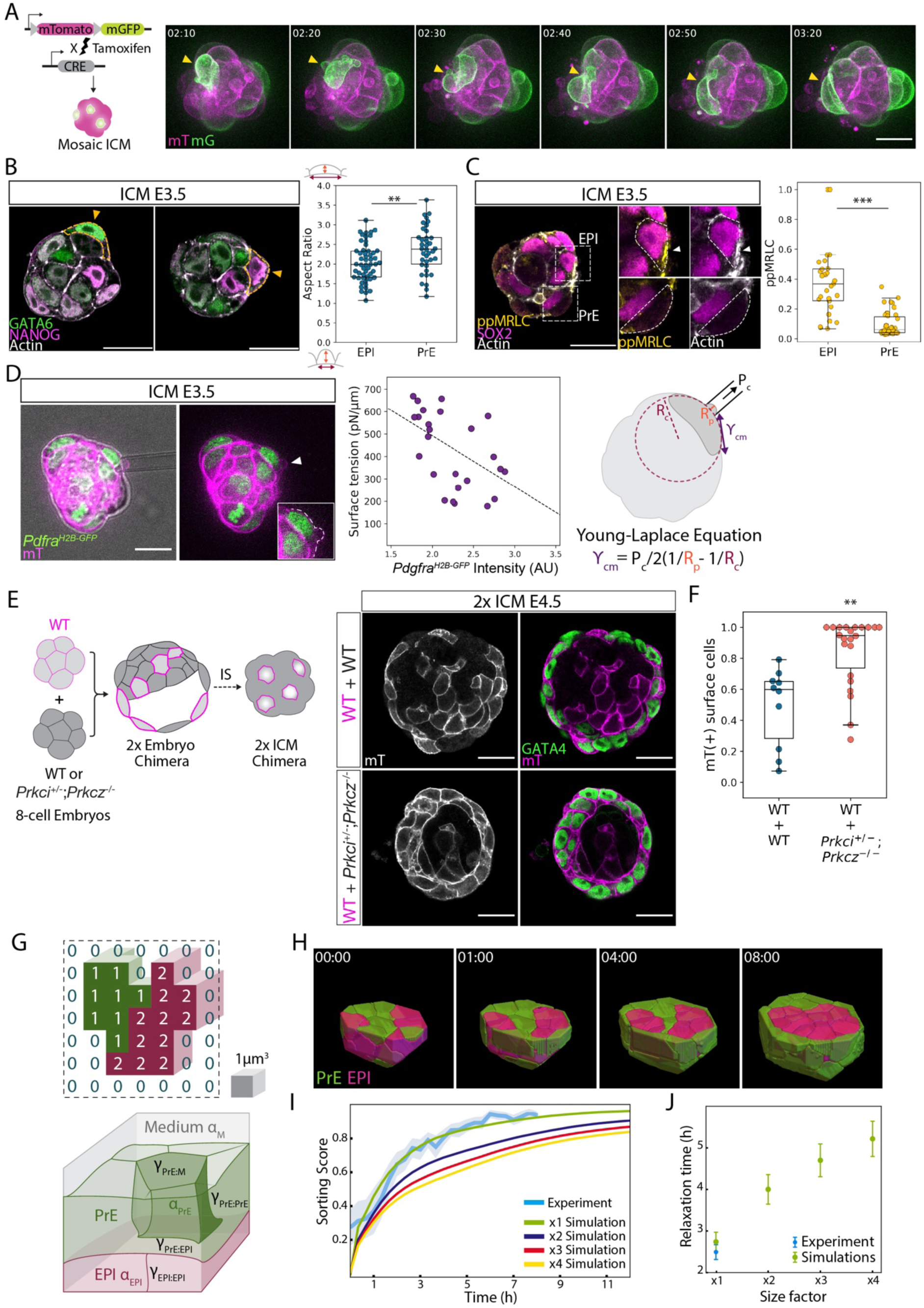
Acquisition of the apical domain decreases surface tension and is sufficient for retaining PrE cells at the fluid interface. A. Schematic representation and time-lapse images of a mosaic-labelled ICM isolated from an *R26CreER*; mTmG blastocyst. Time is indicated as hh:mm, t=00:00 marks stage E3.5+3hrs. Yellow arrowheads, cell shape changes in a surface cell. B. Representative immunofluorescence images of E3.5 ICMs highlighting cell shape among EPI and PrE at the ICM-fluid interface. Analysis of surface cell aspect ratio from E3.5 isolated ICMs to compare EPI and PrE cell shape. n= 53, 38 cells from 16 ICMs. Mann Whitney U test, *p*=6.44e^-03^. C. Immunofluorescence image of an E3.5 isolated ICM showing the distribution of ppMRLC and Actin in EPI (top) and PrE (bottom) cells on the ICM surface, and quantification of normalised ppMRLC distribution at the outer cell cortex. n=29 EPI, 42 PrE cells from 10 ICMs. Mann Whitney U test, p=2.18e^-09^. D. Micropipette aspiration of E3.5 ICMs expressing *Pdgfrα^H2B-GFP^* (green) and membrane td tomato (mT, magenta), and scatter plot of measured surface tension of outer cells versus *Pdgfrα^H2B-GFP^* expression level of the cell. White arrowhead marks the site of cell aspiration and the white dotted line indicates cell surface contour. n= 25 cells from 11 ICMs. Black dotted line, linear regression with Pearson’s R= -0.53, *p*=2.28e^-04^. Interfacial tension is calculated using Young-Laplace equation with γ_cm_, cell-medium interfacial tension, P_c_, aspiration pressure, R_p_, radius of pipette and R_c_, curvature radius of cell surface. E. Schematic for the experimental strategy using chimeric ICMs to test the retention hypothesis, and immunofluorescence images of 2x chimeric ICMs composed of cells from WT+WT combination (top) and WT + *Prkci^+/-^; Prkcz^-/-^*combination (bottom). F. Analysis of the surface retention of WT vs *Prkci^+/-^; Prkcz^-/-^*cells in chimeric ICMs. The plot indicates the proportion of all surface cells that are mT(+) for WT+WT combination and and WT + *Prkci^+/-^; Prkcz^-/-^* combination. n= 10, 22 ICMs for the two groups, respectively. Mann-Whitney U test, *p* = 1.08e^-03^. G. Schematic diagram of a 3D cellular Potts model. The system state is given by a collection of voxels with one of the following three identities – 0 for medium, 1 for PrE and 2 for EPI. Cell-cell and cell-medium interfaces have different tensions γ_PrE:M_ < γ_PrE:PrE_ < γ_EPI:EPI_< γ_PrE:EPI_ < γ_EPI:M_, and the two cell types have different kinetic parameters α_Epi_ < α_PrE_. See Materials and Methods for details. H. Snapshots from the simulation of EPI/PrE sorting in normal-sized isolated ICMs. Poissonian dynamics directly determines the physical time of simulations indicated in hh:mm, with t=00:00 corresponding to E3.5+3hours. I. The time-series of sorting scores of isolated ICMs from experimental live-imaging data shown in Figure 1C, represented as mean SEM, is compared with simulations of isolated ICMs of different size factors. J. Relaxation time 1″ of sorting in isolated ICMs for experimental live-imaging data and simulations of ICMs of different size factors. Scale bars 20μm. *ns, non-significant, * p ≤ 0.05, ** p ≤ 0.01, *** p ≤ 0.001*

To characterise the underlying mechanics generating this PrE cell shape change and difference from EPI cells, we immunostained the ICMs for acto-myosin cytoskeletal elements. Among surface cells, the EPI cell cortex clearly showed higher accumulation of bi-phosphorylated myosin regulatory light chain (ppMRLC) and actin (Figure 2C), suggesting higher acto-myosin activity in EPI cells. Direct measurement of surface tension at the ICM-fluid interface by micropipette aspiration showed that cell-fluid interfacial tension negatively correlates with the *Pdgfrα^H2B-GFP^* intensity (Figure 2D), indicating that EPI cells have higher interfacial tension than PrE cells at the ICM-fluid interface. Furthermore, immunostaining of aPKC isoforms showed their localisation on the contact-free surface of PrE cells but not of EPI cells (Figure S2C,D). These findings suggest that the cell-fluid interfacial tension is reduced at the apical domain in PrE cells, similar to the role of apical polarity in trophectoderm cells of the 16-cell stage embryo (Maître et al. 2016), thereby enabling retention of PrE cells at the ICM surface.

To experimentally test the functional role of the apical polarisation in cell positioning in the ICM, we generated chimeras between fluorescently labelled WT embryos and those lacking aPKC isoforms (Figure 2E). In control chimeras, where fluorescent and non-fluorescent WT embryos are combined, both fluorescent and non-fluorescent cells contributed to the PrE layer at the ICM-fluid interface. However, in aPKC knock-out chimeras, majority of the surface cells were derived from fluorescent WT embryos, whereas non-fluorescent aPKC knock-out cells mostly accumulated inside (Figure 2E,F). Taken together, these experiments demonstrate that apical polarisation is sufficient for retaining PrE cells at the fluid interface due to the lower interfacial tension relative to EPI cells.

### Differential surface tension can sort EPI and PrE cells, but only for the normal-size ICM

Our findings suggest that differential cell-fluid tension may explain EPI/PrE sorting, similar to the mechanism of ICM/TE sorting in the 16-cell stage mouse embryo (Maître et al. 2016). We tested this hypothesis by performing computational simulations of a custom cellular Potts model (CPM) (Glazier and Graner 1993; Graner and Glazier 1992; Hirashima, Rens, and Merks 2017; Savill and Hogeweg 1997; Scianna and Preziosi 2012; Voss-Böhme 2012), which relies on a Poissonian stochastic dynamics instead of the traditional Metropolis scheme (Belousov et al. 2023, *in preparation*, Figure 2G). Our implementation directly introduces physical time into the CPM framework and allows for a fine control of sorting kinetics, thereby enabling the quantification of the time required for sorting in ICMs (see Materials and Methods). To simulate the sorting process, we chose the cell tension parameters from the experimentally observed range of values (see Figure 2D) and found a non-unique set of the remaining parameter values that recapitulate the exponential-like relaxation of the EPI/PrE sorting score observed in our live-imaging experiments with isolated ICMs (Figure 2H, I, and Video S3). To test the robustness of this mechanism against embryo size variation observed *in utero* and with experimental manipulations (Ichikawa et al. 2022; Saiz et al. 2016, 2020), we simulated the sorting dynamics with larger embryos. Notably, whereas the normal-size embryos saturated at a 90% sorting score with a relaxation time of about 2.5 hours, larger embryos took 4-5 hours longer and achieved slightly lower sorting scores (Figure 2I, J, Figure S2E). If surface tension differences are the sole driving mechanism, larger ICMs are thus expected to require a longer time for sorting. Given that the physiological time window for ICM sorting in the developing embryo is approximately 12 hours, with the process commencing around E3.75 and completing by E4.5, the viability and the maintained EPI/PrE proportion of double-size mouse embryos (Saiz et al. 2016, 2020; Tarkowski 1961) suggest additional mechanisms that facilitate the process. Thus, while our simulation shows that differential fluid-cell tension may explain the timely sorting of EPI/PrE cells in normal-size ICMs, this mechanism is not robust to variations that mouse embryos have in developmental progression and embryo size (Fabrèges et al. 2023; Ichikawa et al. 2022). This points to additional mechanisms contributing to EPI/PrE sorting within the ICM that ensure robust patterning of the mouse blastocyst.

### Directed migration of PrE cells depends on actin polymerisation and Rac1

We therefore looked further into cellular dynamics during ICM cell sorting and noted that overall, inner PrE cells show more outward movement in comparison to inner EPI cells (see Figure 1H). To investigate the differential cell dynamics and its underlying mechanisms, however, the small size and spherical geometry of the ICM system limit the interpretation of the analysis, in particular for cells around the centre of the ICM. Visualisation of cell membrane is also necessary to fully characterise cellular dynamics. We thus generated large, mosaic blastocysts using fluorescent reporters marking cell fate with *Pdgfrα^H2B-GFP^*, and membrane with mTmG (Figure 3A, Figure S3A, and Video S4). Live-imaging of these mosaic labelled cells clearly showed distinct cell motility between EPI and PrE. PrE cells migrate towards the fluid-filled cavity with protrusions, while EPI cells remain within the ICM (Figure 3B,C, Figure S3B, see also Video S5, S6), in agreement with the dynamics noted in the ICM culture (Figure 1F,H). Remarkably, PrE cells exhibit a variety of cell shapes, whereas EPI cells remain largely spherical (Figure 3D), and the protrusions of PrE cells are predominantly directed towards the blastocyst cavity (Figure 3E, compare with Figure S3C), indicative of their directed migration. Notably, the directed migration and the length of PrE protrusions, 13.4 μm on average and 18.8 µm at 95^th^ percentile (Figure S3D), are independent of the distance to the cavity interface (Figure 3F).

**Figure 3:**
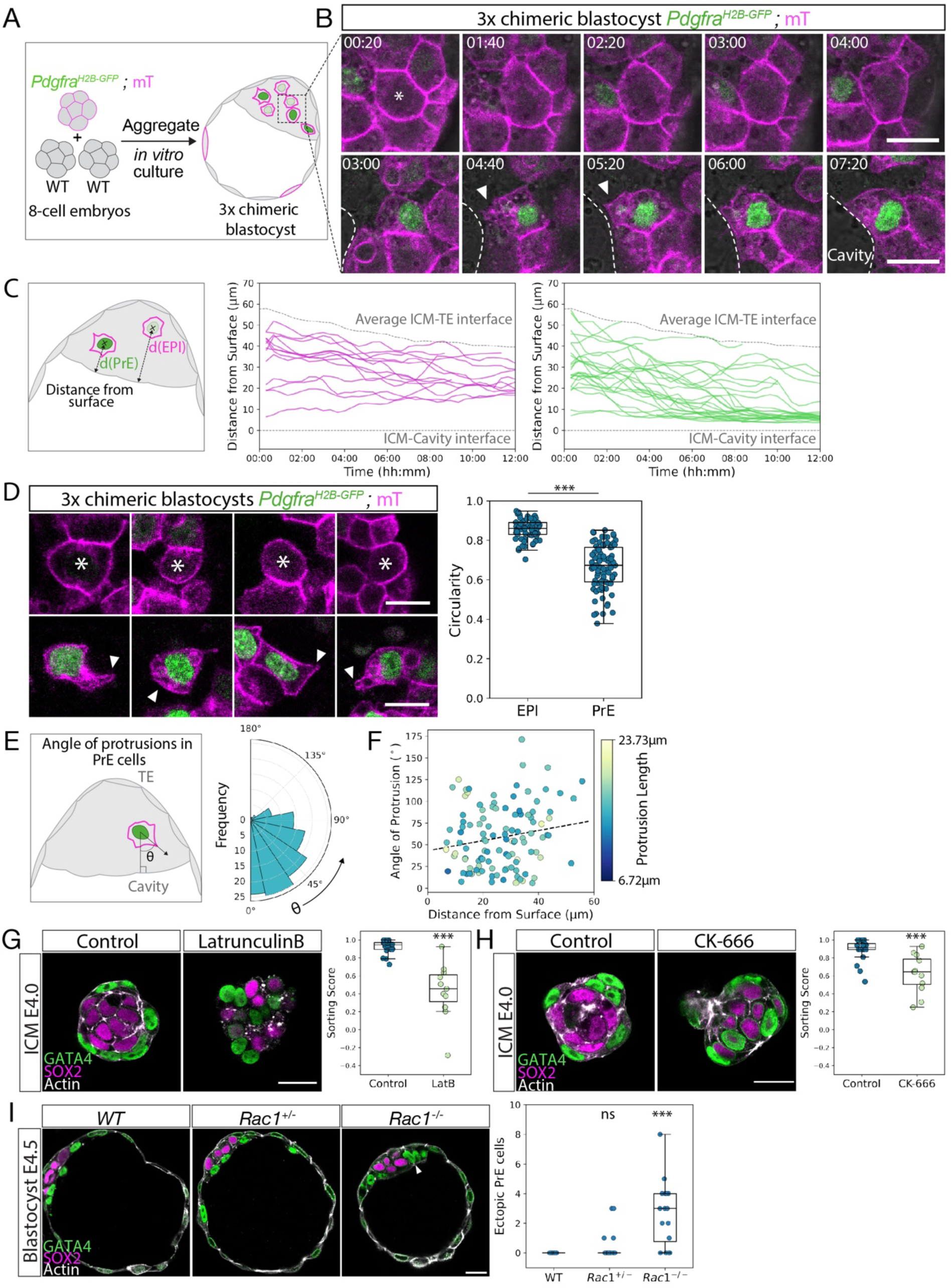
Cell sorting involves active directed migration of PrE cells towards the surface via actin-mediated protrusions. A. Schematic representation of the experimental strategy of aggregating 8-cell stage embryos to generate mosaic labelled cells in large blastocysts to visualise EPI and PrE cell dynamics. B. Time-lapse images of a representative EPI cell (top) and PrE cell (bottom) expressing *Pdgfrα^H2B-GFP^* (green) and membrane td-tomato (magenta) from mosaic labelled blastocysts. White dotted lines mark cavity surface. White asterisk, EPI cell of interest, and white arrowheads, membrane protrusions in PrE cells. Time is indicated as hh:mm, t=00:00 corresponds to start of live-imaging at stage E3.5+3hours. C. Single-cell tracking and analysis of distance of fluorescence-labelled EPI and PrE cells from the cavity surface. Distance is measured from the centre of nucleus/cell to the cavity interface visualised in brightfield. Grey dotted lines mark the average position of ICM-TE interface, and ICM-cavity interface set at d=0. n=14 EPI cells, 31 PrE cells from 13 embryos. D. Representative images and quantification of EPI (top) and PrE (bottom) cell shapes in E3.75 blastocysts. White asterisks, EPI cells of interest. White arrowheads, PrE cell protrusions. Protrusions are defined as membrane deformations that are first extended and then retracted in subsequent timepoints. Mann-Whitney U-test, *p*=7.9e^-22^. n=68 EPI cells, 84 PrE cells. E. Schematic representation and polar histogram of direction of PrE cell protrusions with respect to the cavity. Angle is measured between the segment joining cell nucleus and protrusion tip, and a normal from the nucleus to the cavity interface. n=113 measurements from 12 embryos. Mann-Whitney U-test compared to Figure S2B, *p*=8.41e^-04^. F. Scatter plot of angle of protrusions in PrE cells versus distance of the cell from the cavity. Dots are coloured according to the length of the protrusion as indicated on the colour scale. Dotted line, linear regression. Pearson’s R=0.179, *p*=0.058. n=113 measurements from 12 embryos. G. Representative images of control and latrunculinB-treated E4.0 isolated ICMs and quantification of sorting score. n=20, 11 ICMs for control and latrunculinB-treated ICMs, respectively. Mann-Whitney U test, *p* = 6.16e^-06^ H. Representative images of control and CK-666-treated E4.0 isolated ICMs and quantification of sorting score. n=20, 12 ICMs for control and CK-666-treated ICMs, respectively. Mann-Whitney U test, *p* = 3.69e^-04^ I. Immunofluorescence images of representative WT, *Rac1^+/-^*, and *Rac1^-/-^* E4.5 blastocysts and quantification of number of ectopic PrE cells in WT, *Rac1^+/-^*, and *Rac^-/-^* E4.5 blastocysts. White arrowhead, ectopic PrE cell. n=9, 17, 16 blastocysts respectively. Mann Whitney U test, *p* = 0.133 for comparison between WT and *Rac1^+/-^*, *p*=0.001 for comparison between WT and *Rac1^-/-^*. Scale bars 20μm. *ns, non-significant, * p ≤ 0.05, ** p ≤ 0.01, *** p ≤ 0.001*

To test whether PrE cells actively migrate towards the ICM-cavity interface, we first pharmacologically disrupted actin polymerisation with latrunculin B. This effectively diminished the spatial segregation between EPI and PrE in ICMs (Figure 3G) without compromising cell survival or proliferation (Figure S3E). Secondly, targeted inhibition of actin branching by blocking Arp2/3 activity with CK-666 resulted in failure of PrE cells to reach the ICM surface (Figure 3H). Finally, pharmacological and genetic perturbation of Rac1, a small GTPase essential for active cell migration, led to the failed segregation of PrE cells to the ICM surface as a uniform layer (Figure 3I, Figure S3F, see also Video S7), again without change in the ICM cell number (Figure S3G,H). Collectively, these data show that Rac1 activity and branched actin-mediated protrusions drive directed migration of PrE cells towards the ICM cavity interface during EPI/PrE sorting.

### Apical polarisation in PrE cells is required for directed migration and sorting

To identify what causes the Rac1 activation and branched-actin network in PrE cells, we examined our single-cell gene-expression database for genes differentially expressed between PrE and EPI cells at the beginning of their lineage segregation at E3.5 (Ohnishi et al. 2014). In addition to *Fgfr2*, genes encoding extracellular matrix components such as Laminin α1, Laminin β1, Laminin γ1, and Collagen IV, and factors involved in their synthesis such as Serpinh1 and P4ha2, as well as PKCλ, are specifically expressed in PrE cells (Figure S4A,B; Ohnishi et al. 2014). Immunostaining of the embryo confirmed dense accumulation of laminin and aPKC in PrE cells (Figure 4A,B, FigureS4C). Notably, aPKC is localised near the leading edge of migrating PrE precursors in the E3.75 blastocyst (Figure 4B). To quantitatively analyse the aPKC localisation in PrE cells, we measured the accumulation of aPKC at the sub-cortical region in PrE and EPI cells, which showed that aPKC is differentially localised in PrE cells at the side facing towards the ICM surface (Figure 4C,D). GATA6-expressing PrE cells are more polarised than EPI cells (Figure 4E), independently of cell position within the ICM (Figure 4F), suggesting that PrE cells acquire apical polarity in a cell-autonomous, fate-dependent manner. These findings led us to the hypothesis that the apical polarisation in PrE cells may be functionally linked with their front-rear polarity for directed migration.

**Figure 4:**
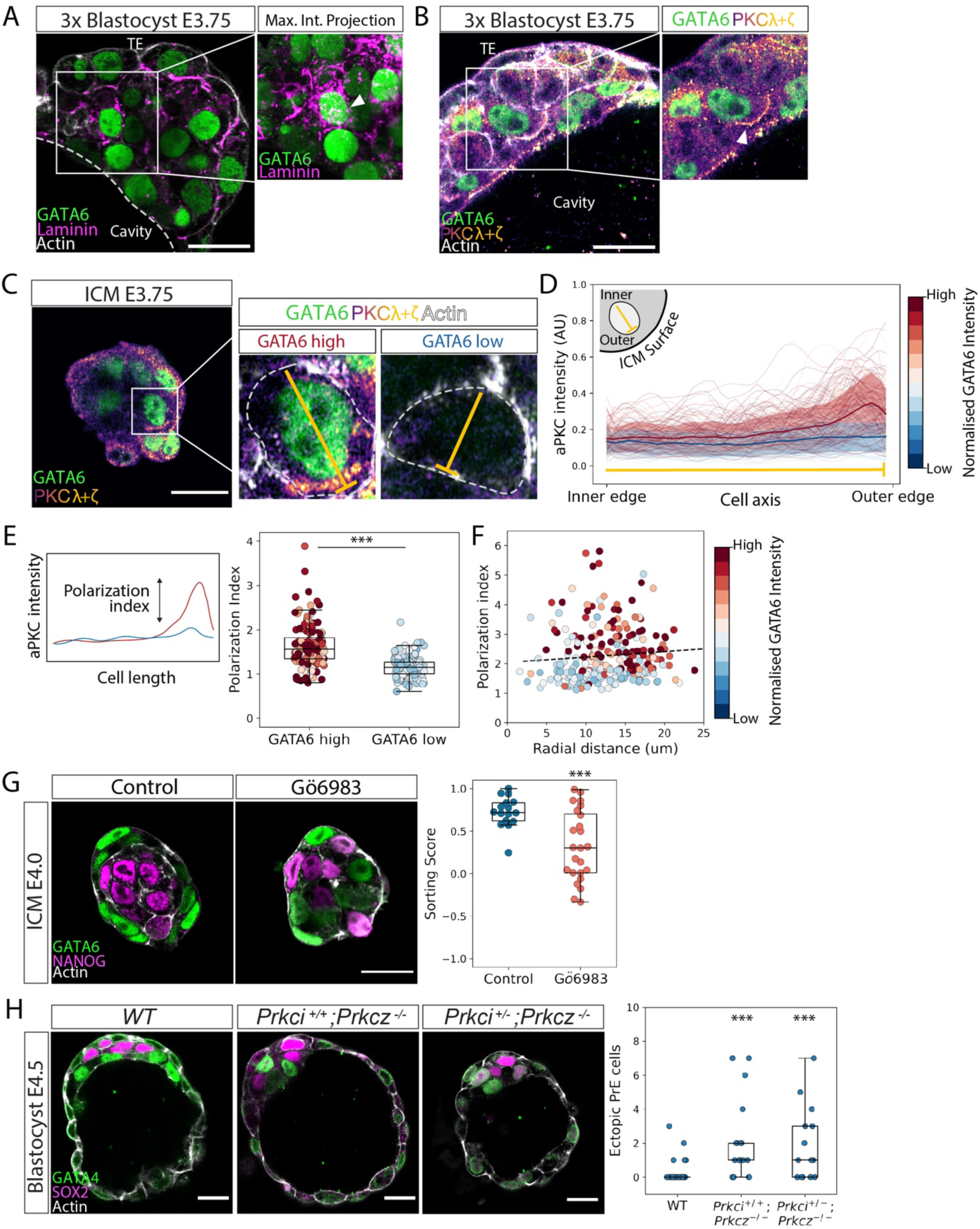
Apical polarisation in PrE cells is required for directed migration and sorting. A. Immunofluorescence image of a 3x blastocyst at stage E3.75 showing laminin distribution around PrE cells. White dotted line, ICM-cavity interface. White arrowhead marks GATA6-expressing nucleus of a PrE cell enriched for laminin expression. B. Immunofluorescence image of a 3x blastocyst at stage E3.75 showing PKCλ+ζ distribution in PrE cells. White arrowhead marks leading edge of a PrE cell with PKCλ+ζ localisation. C. Immunofluorescence image of an E3.75 ICM showing PKCλ+ζ localisation in PrE and EPI cells. White dotted lines mark cell boundaries. Yellow line indicates the line segments from cell inner edge (towards ICM centroid) to cell outer edge (towards ICM-fluid interface) along which fluorescence intensity is measured. D. Line plots for normalised fluorescence intensity of PKCλ+ζ in individual inside cells from E3.75 isolated ICMs. Colour of the line indicates GATA6-expression level of the cell. n=260 cells from 32 ICMs. Each of the thin lines corresponds to measurement from one cell. Bold line and shaded region indicate mean±SD of aPKC intensity for GATA6-high and GATA6-low cells. E. Schematic description of polarisation index. Polarisation index is calculated as the ratio between mean aPKC intensity at 1/4^th^ distance from outer edge and mean aPKC intensity at 1/4^th^ distance from inner edge. Boxplots for comparison of the polarisation index in PrE (GATA6 high) versus EPI cells (GATA6 low). GATA6 expression level is categorised as high or low by thresholding the bimodal distribution of GATA6 fluorescence intensity. Colour of the line indicates GATA6-expression level of the cell. n=136 GATA6-high and 124 GATA6-low cells from 32 ICMs. One-way ANOVA, *p*=6.03e^-20^. F. Scatterplot of polarisation index of cells versus radial distance of the cell from the ICM centroid. Colour of the datapoint indicates GATA6-expression level of the cell. Black dotted line, linear regression with Pearson’s R=0.079, *p*=0.205. n=260 cells from 32 ICMs. G. Immunofluorescence images of control and Gö6983-treated E4.0 isolated ICMs and quantification of sorting score. n=16,24 ICMs for control and Gö6983-treated ICMs respectively. Independent samples t-test, *p* = 8.01e^-04^ H. Immunofluorescence images of representative WT, *Prkci^+/+^;Prkcz^-/-^*, and *Prkci^+/-^;Prkcz^-/-^*E4.5 blastocysts and quantification of number of ectopic PrE cells in E4.5 blastocysts from each group. n= 25, 17, 12 blastocysts for WT, *Prkci^+/+^;Prkcz^-/-^*, and *Prkci^+/-^;Prkcz^-/-^*respectively. Mann-Whitney U test, *p* = 2.43e^-04^, 6.36e^-04^ Scale bars 20μm. *ns, non-significant, * p ≤ 0.05, ** p ≤ 0.01, *** p ≤ 0.001*

To test the functional role of aPKC in PrE cell migration, we first inhibited the activity of aPKC in the ICM with Gö6983, which disrupted sorting and patterning in agreement with a previous report (Figure 4G, Figure S4D, see also Video S8; Saiz et al. 2013). Furthermore, combined genetic knock-outs of aPKC isoforms, *Prkci^+/+^; Prkcz^-/-^* and *Prkci^+/-^; Prkcz^-/-^*, resulted in smaller ICMs (Figure S4E) with failed segregation of PrE to the cavity surface (Figure 4H, and Video S9), indicating that functional apical polarity is necessary for directed migration and sorting of PrE cells. Taken together, early in differentiation and within the ICM, PrE cells acquire the apical polarity that is required for directed migration and sorting to the ICM-cavity interface.

### ECM deposited by PrE cells builds a tissue-level gradient in the ICM and guides directional PrE cell migration

Thus far, our findings showed that acquisition of apical polarity is required and sufficient for PrE cell migration and surface retention, respectively. However, it is unclear what directs PrE cells within the ICM tissue to migrate towards the ICM surface, in particular towards the ICM fluid interface in the blastocyst. PrE cells near the cavity may be trapped to the surface when protrusions reach the fluid interface, though this mechanism *per se* cannot explain the directed protrusions of PrE cells deeper than 20μm from the cavity (see Figure 3E,F). We reasoned that this surface trapping of PrE cells near the cavity, however, may break the tissue-level symmetry with respect to the distribution of salt-and-pepper PrE and EPI cells, hence the distribution of ECMs secreted by PrE cells, which can in turn guide the PrE cell migration at the tissue scale.

This hypothesis predicts that the distribution of ECM components, while uniform before sorting, progressively form a gradient across the ICM tissue, highest around the cells facing the blastocyst cavity. To test this prediction, we immunostained enlarged blastocysts for pan-laminin to gain higher spatial resolution and quantified its distribution across the ICM. This revealed that the uniform distribution of laminin at E3.5 indeed changes to build a gradient at E3.75 when EPI/PrE sorting takes place (Figure 5A,B). To determine whether ECM is functionally active and whether cell-matrix adhesion is present in PrE cells, we coimmunostained active integrinβ1 and pan-laminin (Figure 5C, Figure S5A). This showed a large fraction of active integrinβ1 colocalised with laminin, strongly suggesting the presence of active cell-matrix adhesion. These findings support the model in which migration of the PrE cells near the cavity surface break symmetry in ECM distribution (Figure S5B) and build a gradient across the ICM tissue, which can then guide other PrE cells to migrate towards the cavity surface (Figure 5D). This is also in line with our earlier findings that integrinβ1 and laminin γ1 are required for proper PrE segregation to the ICM surface (Kim, Sorokin, and Hiiragi 2022).

**Figure 5:**
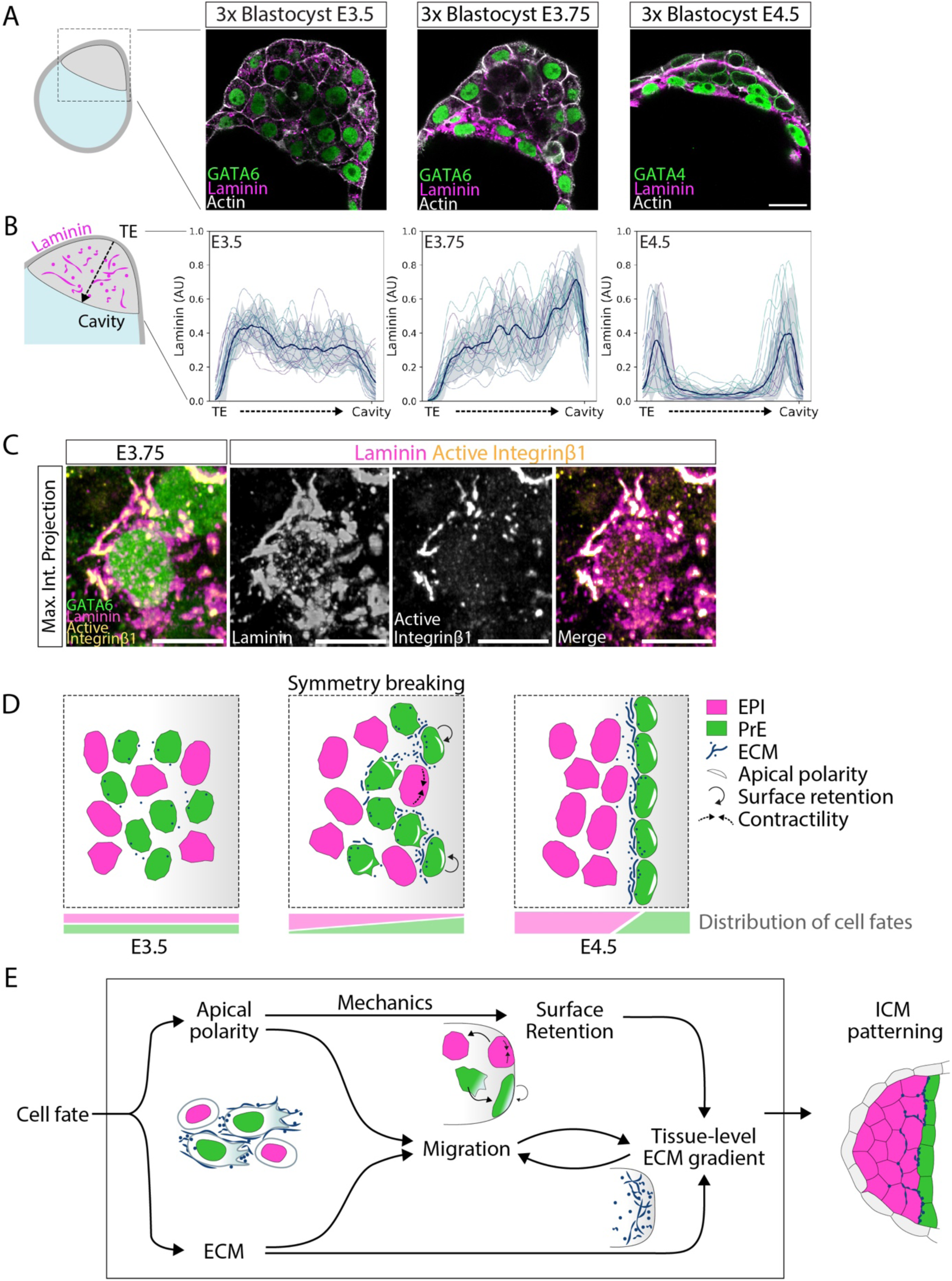
Extracellular matrix deposited in the ICM guides PrE cells towards the cavity surface. A. Immunofluorescence images of ICMs in 3x blastocysts at stages E3.5, E3.75 and E4.5 stained for GATA4, pan-laminin, and actin. B. Quantification of total pan-laminin distribution from the ICM-TE interface to the ICM cavity interface in maximum intensity projections of the blastocysts from (A). The solid line and shaded region indicate mean±SD of the laminin fluorescence intensity. Line plots were constructed with data from n=5, 6, 5 embryos for the different stages respectively. Lines of the same colour correspond to measurements from the same embryo at respective stages. C. Maximum intensity projections of a representative immunofluorescence image indicating colocalisation of laminin and active integrinb1 (9EG7) in PrE cells in the ICM in 3x blastocysts at stage E3.75. Manders’ coefficients for colocalised fractions are: Laminin overlapping Itgb1 = 0.185, and Itgb1 overlapping Laminin = 0.891. D. Schematic representation to explain EPI-PrE fate segregation. Until stage E3.5, there is negligible asymmetry in ICM composition. Around stage E3.75, PrE cells in the ICM acquire hallmarks of apical polarity, and begin to express and secrete ECM components. Polarisation of PrE cells has two major consequences – PrE cells already at the cavity undergo surface retention, as apolar EPI cells have higher surface tension and move inwards. Secondly, inner PrE cells extend cell protrusions that facilitate their migration towards the cavity, as collectively guided by the higher distribution of ECM, to result in pattern formation at the tissue level. E. Multi-scale feedback model of tissue level patterning between cell polarisation, mechanics, cell migration and ECM deposition triggered by the establishment of cell fates to explain blastocyst patterning Scale bars 20μm.

Taken together, these data led us to a mechanistic model of EPI/PrE sorting that integrates cell fate, polarity, mechanics and tissue-scale positional information (Figure 5E). First, within the ICM tissue, salt-and-pepper-distributed PrE cells acquire apical polarity that induces cell protrusive and migratory activity. Protrusions from PrE cells near the cavity reach the fluid interface and induce their retention at the surface, which shifts the balance of PrE cell distribution, and thereby that of their secreted ECM. The progressively increasing asymmetry in the ECM distribution can guide other PrE cells to migrate towards the cavity, which in turn contribute to the emerging tissue-level ECM gradient, effectively enabling collective cell migration towards the surface – which we term “breadcrumb navigation” (Figure 5D). This multi-scale feedback model explains tissue-level symmetry breaking and dynamic pattern emergence within an initially equivalent population of cells (Figure 5E).

### The fixed proportion of EPI/PrE cells without cell fate plasticity challenges precision in ICM patterning

While this feedback model may explain dynamic EPI-PrE cell segregation and pattern emergence, we sought to understand how this mechanism is linked with cell fate specification. Cell lineage, division pattern and gene expression pattern are all variable among embryos in pre-implantation mouse development, and in such a system, feedback of cell positional information to cell fate specification could ensure robust patterning (Chan et al. 2019; Kim, Korotkevich, and Hiiragi 2018; Korotkevich et al. 2017; Maître et al. 2016). In line with this model, earlier studies (Meilhac et al. 2009; Plusa et al. 2008; Wigger et al. 2017) proposed position-dependent PrE fate specification, in which cells on the cavity surface are induced to differentiate into PrE. However, this is incompatible with another notion that the proportion of EPI/PrE cells is fixed according to the gene regulatory network between GATA6, NANOG and FGF signalling activity (Bessonnard et al. 2014; Kang et al. 2013; Krawchuk et al. 2013; Saiz et al. 2016, 2020; Schrode et al. 2014; Yamanaka, Lanner, and Rossant 2010).

First, we analysed the proportion of EPI/PrE cells in blastocysts and ICMs experimentally isolated from blastocysts, and found it indeed constant (Figure 6A) with PrE proportion 0.606 ± 0.08 (n=106 embryos), in agreement with earlier studies (Saiz et al. 2016, 2020). Next, cell lineage tracking with fate marker showed highly limited contribution of position-dependent fate-switching during the EPI/PrE sorting; only 3 of 181 cells differentiate to PrE by increasing the expression of *Pdgfrα^H2B-GFP^* on the cavity surface, whereas 2 of 93 cells differentiate to EPI by decreasing the *Pdgfrα_H2B-GFP_* signal inside the ICM (Figure 6B). These findings suggest that the fate and proportion of EPI/PrE cells are fixed in E3.5-4.5 blastocysts.

**Figure 6:**
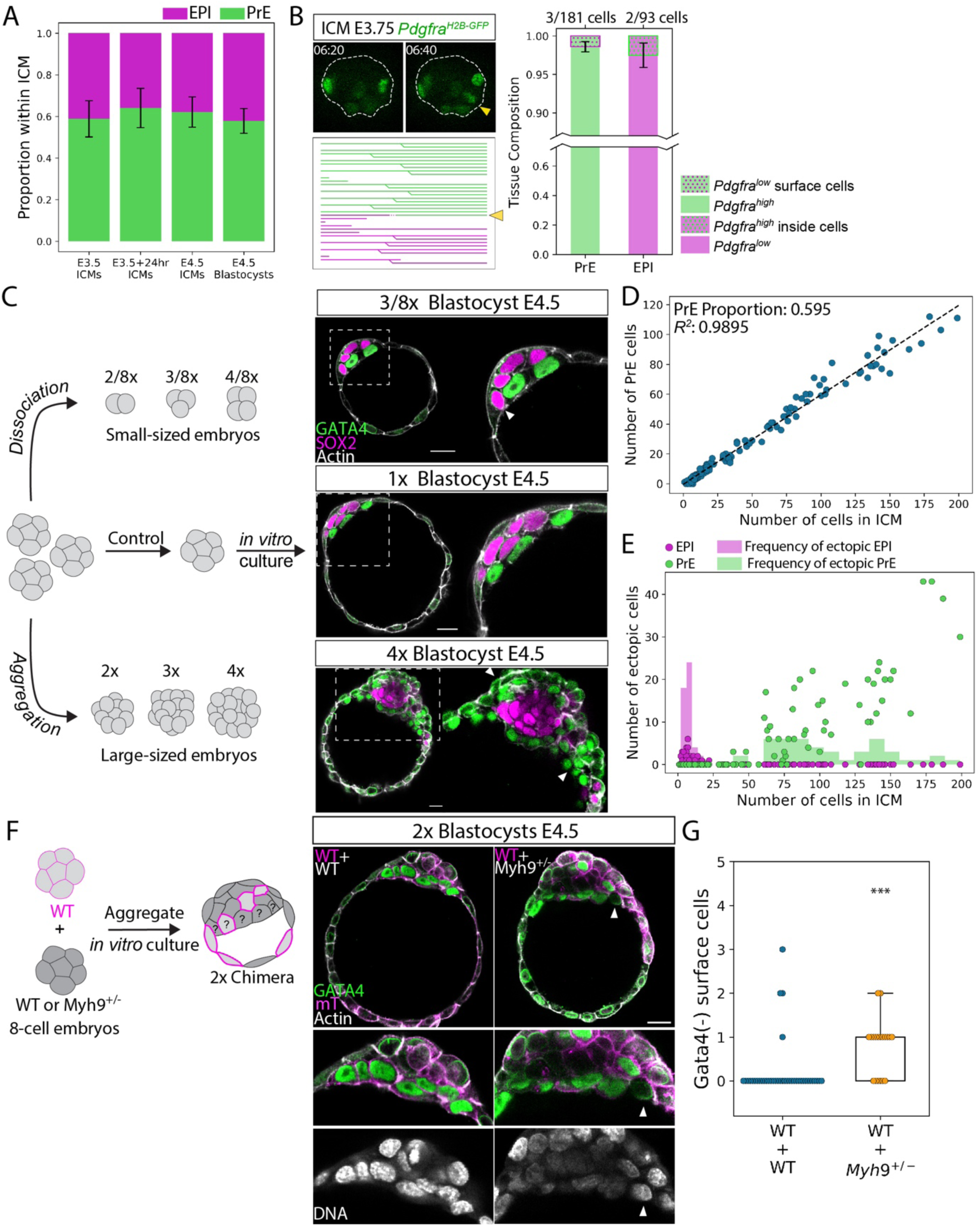
The fixed proportion of EPI/PrE cells without cell fate switching challenges precision in ICM patterning. A. Proportion of cell fates in the ICM in embryos under different conditions during development (PrE: green, and EPI: magenta). One-way ANOVA, *p*=0.085. n=19, 29, 21, 32 embryos for the different groups, respectively. Mean PrE proportion = 0.605±0.78. B. Limited contribution of position sensing and cell fate switching in E3.75 ICMs to final patterning of the ICMs. Consecutive time-lapse images from isolated ICMs expressing *Pdgfrα^H2B-GFP^* and the corresponding lineage tree for the ICM. White dotted line marks ICM boundary. t=00:00 corresponds to start of live-imaging at stage E3.5+3hours, following completion of immunosurgery. Lineage tree of an isolated ICM from single-cell tracking in Figure 1C,D. Yellow arrowhead marks an inside cell that increases *Pdgfrα^H2B-GFP^* expression after moving to the surface, and its representation on the lineage map. Stacked bar plots for quantification of the frequency of position sensing and fate switching contributing to the final EPI and PrE cell fates. C. Immunofluorescence images of size-manipulated blastocysts indicating defects in ICM patterning at stage E4.5. White arrowheads mark ectopic EPI cells in smaller blastocysts and ectopic PrE cells in larger blastocysts. D. Number of PrE cells plotted as a function of total number of cells in the ICM in E4.5 size manipulated blastocysts. Dotted line, linear regression with Pearson’s R=0.991, *p*=1.18e^-134^. PrE proportion = 0.595 ± 0.005. n = 29 embryos for 2/8x, 26 embryos for 3/8x, 17 embryos for 4/8x, 29 embryos for 1x, 19 embryos for 2x, 24 embryos for 3x, 10 embryos for 4x size ratios. E. Quantification of ectopic EPI and PrE cells in size-manipulated E4.5 blastocysts. The number of ectopic cells is plotted as a function of total number of cells in the ICM. Magenta, ectopic EPI cells and green, ectopic PrE cells. Frequency of the occurrence of ectopic cells with increasing ICM cell number is indicated as shaded histogram. F. Schematic representation of chimera experiments to test feedback in the ICM between cell fate and position. Immunofluorescence images of 2x chimeric E4.5 blastocysts composed of cells from WT+WT combination (left column) and WT + *Myh9^+/-^* combination (right column). White arrowhead marks GATA4-negative cells on the ICM surface. G. Quantification of number of GATA4-negative cells on the cavity surface in WT+WT combination and WT + *Myh9^+/-^* combination of E4.5 chimeric blastocysts. n= 40, 19 embryos for the two groups, respectively. Mann-Whitney U test, *p* = 2.82e^-05^ Scale bars 20μm. *ns, non-significant, * p ≤ 0.05, ** p ≤ 0.01, *** p ≤ 0.001*

To distinguish the presence or absence of position-dependent cell-fate plasticity, we challenged the system by manipulating embryo size 4-fold larger or smaller. These major changes in ICM cell number (Figure S6A) lead to corresponding differences in the ICM interface/volume ratio (Figure S6B, C), which, in the absence of position-dependent cell-fate plasticity, would result in failure to fit one layer of PrE cells on the ICM surface (Figure 6C, see also Video S9). In agreement with the notion of fixed EPI/PrE proportion, we found that, despite the wide range of variability in the number of ICM cells in larger or smaller embryos, the proportion of PrE remained constant at 0.595±0.005 (n=144; Figure 6D). Remarkably, in larger embryos, we observed ectopic PrE cells within the ICM, and conversely, in smaller embryos, ectopic EPI cells at the ICM-fluid interface (Figure 6C,E, see also Video S9). These findings support the lack of cell fate plasticity at this stage, and the lack of feedback from cell position to fate specification.

To unequivocally demonstrate the presence or absence of cell fate plasticity at this stage, we further challenged the system by generating chimeric blastocysts using embryos heterozygous for *Myh9,* which encodes the myosin heavy chain (*Myh9^+/-^,* Figure 6F). The chimeras between WT and *Myh9^+/-^* embryos would force some EPI cells derived from *Myh9^+/-^* embryos to be located on the ICM surface as they have relatively lower cortical tension. Importantly, *Myh9^+/-^* embryos form blastocysts with ICM cell number and EPI/PrE proportion comparable to WT (Figure S6D, see also Video S10). Without position-dependent cell fate plasticity, these cells would not change fate to PrE despite being on the surface. Control chimeras form a precise ICM pattern at the E4.5 stage without ectopic cells (Figure 6G). In contrast, chimeras with *Myh9^+/-^* embryos show ectopic cells on the ICM surface that do not change fate to PrE, thus experimentally supporting the model that EPI/PrE cell fate and their proportions are fixed without plasticity during ICM patterning in late blastocysts.

### The fixed proportion of EPI/PrE cells is optimal for the specific ICM size and geometry across mammals

The fixed EPI/PrE proportion and the lack of plasticity would present a challenge for early mammalian embryos to achieve precision in blastocyst patterning, as cell numbers, lineage, and embryo geometry are variable among embryos during pre-implantation development. We therefore asked how robust patterning is ensured in the absence of cell fate plasticity. Not only the number of cells but also their shape varies in each embryo. Thus, the variability of cell shape defines the range of cell numbers with which an embryo with a given geometry and fixed EPI/PrE proportion can achieve precise patterning, covering the bulk of EPI cells with an intact PrE monolayer. To estimate this range and examine its distribution across embryos of various sizes, we characterized the *in vivo* geometry of the tissue and individual cells from immunostained blastocysts. Specifically, we approximated the ICM shape as a combination of two spherical caps, corresponding to the ICM-trophectoderm interface, and the ICM-cavity interface, and thus obtained estimates of the ICM-cavity interface area A_Interface_ (Figure 7A). Next, we measured the PrE cell apical areas at the cavity surface, and determined the 10^th^ and 90^th^ percentiles of the cell area q_10%_=157μm^2^ and q_90%_=376μm^2^ respectively (Figure S7A). Given the fixed proportion f=0.6 of PrE cells, we calculated the corresponding range of the total PrE area A_PrE_ = [f n q_10%_, f n q_90%_]. This range is bound by the marginal sizes of a hypothetical monolayer formed by the PrE cells in an embryo with a total cell count n, because the variability in single-cell apical areas gives rise to an interval of possible values that a total PrE area may have (Figure 7A). By comparing A_Interface_ and A_PrE_, we predict the presence of gaps or multilayered regions for different ICM sizes: if A_Interface_ is larger than the maximal bound of A_PrE_, we expect a gap in the PrE monolayer with EPI cells exposed to the interface, whereas if A_Interface_ is smaller than the minimal bound of A_PrE_, superfluous PrE cells would be located inside the ICM thus forming a PrE multilayer.

**Figure 7:**
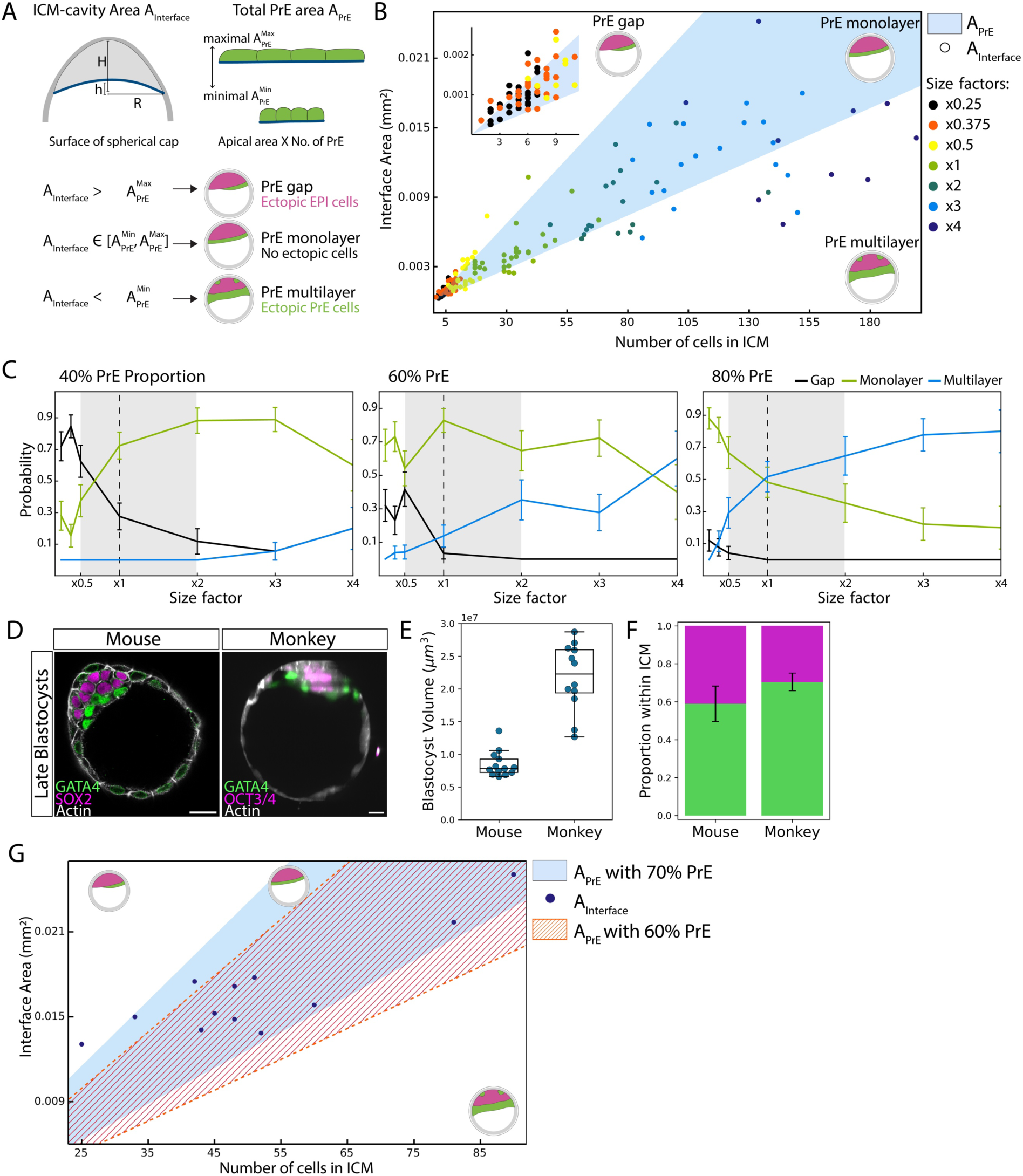
The fixed proportion of EPI/PrE cells is optimal for the specific embryo size and ICM geometry. A. Schematic representation of estimation of ICM-cavity interface area A_Interface_ and total PrE area A_PrE_ in blastocysts. A_Interface_ is calculated from fitting two spherical caps to the ICM geometry, using empirical measurements of ICM base radius R, major height H and minor height h. A_PrE_ is calculated as the product of PrE cell apical area and the number of PrE cells in the ICM. Variability in PrE cell stretching gives rise to an interval of maximal and minimal A_PrE_. B. Scatter plot depicting A_Interface_ as a function of total number of cells in the ICM for size manipulated E4.5 embryos. Blue shaded region corresponds to the maximal and minimal bounds of A_PrE_. Colour of the dots indicates size ratio of the embryos. Inset, zoomed-in region for x0.25 and x0.375 size ratio. C. Estimated probabilities of forming PrE gap, PrE monolayer and PrE multilayer with embryo size for ICM composition of 40% PrE, 60% PrE and 80% PrE. Grey shaded region indicates natural variability in embryo size between size ratios x0.5 and x2. D. Representative immunostaining images of E4.5 mouse blastocysts and monkey blastocysts 7-8 days post-ICSI (intracytoplasmic sperm injection). E. Quantification of blastocyst volume in mouse and monkey blastocysts. n=13, 12 embryos for mouse and monkey, respectively. F. Quantification of EPI/PrE proportion within the ICM for mouse and monkey blastocysts, plotted as mean ± SD. Mean PrE proportion for mouse embryos, 0.59±0.09 and mean PrE proportion for monkey embryos, 0.70±0.05. G. Scatter plot depicting A_Interface_ as a function of total number of cells in the ICM for monkey blastocysts 7-8 days post-ICSI. Blue shaded region corresponds to the maximal and minimal bounds of A_PrE_, assuming 70% PrE proportion within the ICM. Orange dashed region corresponds to the maximal and minimal bounds of A_PrE_, assuming 60% PrE proportion within the ICM. sScale bars 20μm.

Our measurements showed that the majority of normal-sized embryos have ICM-cavity interface areas within the range that PrE cells could cover, thereby enabling formation of a PrE monolayer (Figure 7B, n=29 embryos). Furthermore, by counting the frequency of embryos inside (outside) the region A_Interface_, we estimated the probability of observing a monolayer (gap / multilayer), and found that gap formation is more likely to occur in smaller embryos, and multilayers more likely in larger embryos (Figure 7B, inset; see Figure 6C,E). We then compared the probability of PrE monolayer formation across embryos of various sizes, and found it highest in the normal-size embryo (Figure S7B, 82.5%). Further, the probability of monolayer formation was higher than that of multilayers or gaps for embryos across the four fold size difference (from double to half size), indicating the robustness of precise ICM pattern formation against natural variability of embryo size (Figure 7C). However, when the probability is calculated for the scenario where the ICM is composed of 40% or 80% PrE, the likelihood of forming gaps in the PrE layer is higher in 40% PrE and that of multilayer formation is higher in 80% PrE (Figure 7C). Notably, the probability of monolayer formation without a gap or multilayer is highest with 60% PrE for the mouse embryo for a range from double to half size, suggesting that this fixed EPI/PrE proportion is optimal for patterning the mouse ICM given its size and geometry.

Other mammalian species have different embryo sizes and proportions of EPI/PrE in the ICM (Kuijk et al. 2012; Piliszek et al. 2016; Roode et al. 2012). Our findings, which suggest an optimal proportion of EPI/PrE for a specific embryo size and geometry, therefore raise a question whether different mammalian species have distinct optimal proportions according to their respective sizes and geometries. We tested this prediction by analysing monkey blastocysts. Monkey embryos are larger in size than mouse embryos (Figure 7D,E, see also Video S12), and the ICM has a higher proportion of PrE cells (70%, n=12; Figure 7F). Notably, measurements of cell and ICM geometry show that the observed monkey blastocysts have ICM-fluid interfacial areas within the range that PrE areas could cover when the increased 70% proportion of PrE cells is taken into account (Figure 7G). However, with a 60% PrE ratio–as in mouse embryos–the hypothetic area of PrE cells in monkey embryos would decrease substantially below the observed values (Figure 7G), in line with the notion that an optimal proportion of the ICM cell fates is species-specific. Together, these data suggest that the optimal EPI/PrE ratio may adapt to embryo size and geometry across different species.

## DISCUSSION

Overall, this study addresses the challenge and mechanism for robust ICM pattern formation in mammalian blastocysts. We find that the tissue-level symmetry within the ICM is first broken by retention of PrE cells at the fluid-interface by differential surface tension. This builds a tissue-wide ECM gradient deposited by PrE cells, which guides active migration of PrE cells driven by their acquisition of apical polarity. Despite the fixed proportion of EPI/PrE cells, patterning is robust against naturally variable sizes of the embryo because this proportion is species-specific and optimal for embryo size and geometry.

Although cell fate specification (Bessonnard et al. 2014; Chazaud et al. 2006; Frankenberg et al. 2011; Kang et al. 2013; Kang, Garg, and Hadjantonakis 2017; Krawchuk et al. 2013; Molotkov et al. 2017; Ohnishi et al. 2014; Schrode et al. 2014; Xenopoulos et al. 2015; Yamanaka et al. 2010) and sorting in the ICM (Meilhac et al. 2009; Plusa et al. 2008; Saiz et al. 2013; Wigger et al. 2017; Yanagida et al. 2022) have been studied, they were not combined to measure fate-specific dynamics underlying cell sorting. PrE-specific cell surface fluctuations and differential cell fluidity were recently shown to be sufficient for sorting cell aggregates (Yanagida et al. 2022), though how these properties arise only in PrE cells, and how this could pattern the blastocyst ICM with a specific *in vivo* geometry remained elusive. Here, we report for the first time, that PrE cells undergo Rac1-dependent active migration. Furthermore, we find autonomous polarisation of PrE cells within the ICM, which drives the formation of actin-based protrusions for cell migration, corroborating a functional link between polarity and cell sorting, in agreement with previous findings (Gerbe et al. 2008; Saiz et al. 2013). While apical polarisation has thus far been detected in PrE cells only after their sorting to the fluid interface (Gerbe et al. 2008; Saiz et al. 2013; Yang et al. 2002), here we characterise the asymmetric cortical localisation of aPKC in PrE cells in the salt-and-pepper ICM. This apical polarisation is atypical for two reasons: first, the apical domain usually forms only at the cell-fluid interface (Akhtar and Streuli 2013; Bryant et al. 2010; Korotkevich et al. 2017), but here its formation within the salt-and-pepper cell aggregates was necessary for directed active migration. Second, epithelialisation is typically associated with stabilisation during collective cell migration (Durdu et al. 2014; Friedl and Gilmour 2009), whereas here the apical domain is linked with protrusion and mesenchyme-like motility, which may be due to the immaturity of the apical polarity within the ICM tissue.

PrE cell migration within the ICM is a collective behaviour. Notably, however, this does not require direct alignment of migratory cells, with directional guidance potentially provided by the tissue-level gradient of deposited ECMs, the “breadcrumb navigation”. Cell-ECM interactions can govern aspects of cell migration, as cells *in vivo* enhance their migratory capacity by secreting laminin (Sánchez-Sánchez et al. 2017), and cell-matrix interactions enable cell sensing of a stiffness gradient for durotaxis (Shellard and Mayor 2021). Within the ICM, the ECM accumulates more towards the cavity-interface as EPI/PrE sorting begins. We propose that this gradient is formed by PrE cells themselves, as the initial tissue-scale symmetry is broken by the retention of PrE cells located close enough to the fluid-interface, and whose protrusions reach the surface and drive cell sorting by differential surface contractility. This mechanism can self-organise directed collective cell migration for a certain range in space and time.

Cell sorting at the fluid-interface that can be sufficiently explained by relative differences in surface tension is reminiscent of the mechanism sorting the inside and outside cells in 16-cell embryos (Maître et al. 2016). Enrichment of aPKC at the apical cortex in blastomeres antagonises myosin phosphorylation, which can explain decreased interfacial tension in PrE cells and their retention at the cavity interface. This mechanism couples cell fate and position, ensuring robust patterning, and interestingly, is conserved across two consecutive lineage segregation events in pre-implantation mouse development.

Dynamic mechanisms generating patterns within initially equivalent cell populations described here may be widespread among undifferentiated or stem cell populations, in which stochastically variable gene expression is evident before lineage segregation (Hu et al. 1997; Klein and Simons 2011; Martinez Arias and Brickman 2011; Miao et al. 2023; Simons and Clevers 2011). In such systems, gene regulatory networks or signalling from the niche may drive formation of distinct cell types at a certain ratio, first in a salt-and-pepper pattern, that is subsequently sorted to form a particular pattern within a specific geometrical context. This may present a challenge to developing or homeostatic systems, as they need to accommodate a certain proportion of cell types into varying geometries and environments. Our findings in this study have implications based on the relationship between the ratio of cell types and the geometric properties of the tissue: PrE cells must form a monolayer on the surface of the ICM of varying size and shape, while keeping a fixed PrE/EPI ratio within the ICM. Thus, patterning precision is not always compatible with scaling in tissues. We propose that in such a case, robustness in tissue patterning may be ensured by selecting optimal parameter sets in space and time – in the case of the blastocyst patterning, cell fate proportions, cellular dynamics, duration of sorting, tissue size, and geometry may co-adapt on evolutionary timescales, to be robust against a certain degree of variabilities. Further investigations into mechanisms that enable coordination of these parameter changes will be valuable to gain insights into robustness of embryo development and evolution.

## Acknowledgements

We are grateful to the members of the Hiiragi laboratory and Nicoletta Petridou for discussions and comments on the manuscript; Chii Jou Chan and Esther Kim for help with adapting the culture of isolated ICMs, Dimitri Fabrèges and Anniek Stokkermans for assistance with image analysis; Ramona Bloehs, Stefanie Friese, and Lidia Pérez for their technical support. We thank members of the Tsukiyama group for the animal care with monkeys, in particular Hideaki Tsuchiya and Masataka Nakaya. We thank the EMBL animal facility (LAR) and Animal Care at the Hubrecht Institute for their support. T.I. was supported by the JSPS Overseas Research Fellowship. The Hiiragi laboratory was supported by the EMBL, and currently by the Hubrecht Institute, the European Research Council (ERC Advanced Grant “SelforganisingEmbryo”, grant agreement 742732; ERC Advanced Grant “COORDINATION”, grant agreement 101055287), Stichting LSH-TKI (LSHM21020) and JSPS KAKENHI grant numbers JP21H05038 and JP22H05166. The Erzberger laboratory is supported by the EMBL.

## Author Contributions

Conceptualization, A.E., T.H.; Methodology, P.M., R.B., F.G., A.E., T.H.; Software Programming, R.B.; Validation, P.M., R.B.; Formal Analysis, P.M.; Investigation, P.M., T.I., C.I.; Resources, T.T., T.H.; Data Curation, P.M.; T.I.; Writing – Original Draft, P.M., T.H.; Writing – Review & Editing, P.M., R.B., T.I., F.G., A.E., T.H.; Visualization, P.M., R.B., F.G.; A.E.; Supervision, A.E., T.H.; Project Administration, T.H.; Funding Acquisition, T.H.

## MATERIALS AND METHODS

### Mouse work

All mouse-related animal work was performed in the Laboratory Animal Resources (LAR) Facility at European Molecular Biology Laboratory (EMBL) with permission from the Institutional Animal Care and Use Committee (IACUC) overseeing the operation (IACUC number TH11 00 11), and at the Animal Facility at the Hubrecht Institute. LAR facilities operate according to Federation for Laboratory Animal Science Associations (FELASA) guidelines and recommendations. At the Hubrecht animal facility, mice were housed according to institutional guidelines, and procedures were performed in compliance with Standards for Care and Use of Laboratory Animals with approval from the Hubrecht Institute ethical review board. Animal experiments were approved by the Animal Experimentation Committee (DEC) of the Royal Netherlands Academy of Arts and Sciences. All experimental mice were maintained in specific pathogen-free (SPF) conditions on a 12–12 hour light-dark cycle and used from 8 weeks of age.

### Monkey work

Monkey animal work was performed with female cynomolgus monkeys (*Macaca fascicularis*), of ages ranging between 6 to 11 years. The animals were maintained on a 12-12 hours light dark cycle. Each animal was fed 20 g/kg of body weight of commercial pellet monkey chow (CMK-1; CLEA Japan) in the morning, supplemented with 20–50 g of sweet potato in the afternoon. Water was provided ad libitum. Animals were housed with temperature and humidity maintained at 25 ± 2°C and 50 ± 5%, respectively. The animal experiments were appropriately performed by following the Animal Research: Reporting in Vivo Experiments (ARRIVE) guidelines developed by the National Centre for the Replacement, Refinement & Reduction of Animals in Research (NC3Rs), and also by following “The Act on Welfare and Management of Animals” from Ministry of the Environment, “Fundamental Guidelines for Proper Conduct of Animal Experiment and Related Activities in Academic Research Institutions” under the jurisdiction of the Ministry of Education, Culture, Sports, Science and Technology, and “Guidelines for Proper Conduct of Animal Experiments” from Science Council of Japan. All animal experimental procedures were approved by the Animal Care and Use Committee of Shiga University of Medical Science (approval number: 2021-10-4).

### Mouse lines and genotyping

The following mouse lines were used in this study: C57BL/6xC3H F1 hybrid as wild-type, mTmG (Muzumdar et al. 2007), *Pdgfr⍺^H2B-GFP^* (Hamilton et al. 2003), Prkci^tm1.1Kido^, Prkcz^tm1.1Cda^ (Hirate et al. 2013), R26CreER (Badea et al. 2003), R26-H2B-mCherry (Abe et al. 2011), *Myh9*^tm5Rsad^ (Jacobelli et al. 2010), *Rac1^flox/flox^* (Walmsley et al. 2003), and ZP3-Cre (Lewandoski, Wassarman, and Martin 1997). To generate *Myh9^+/-^*animals, *Myh9^floxed/floxed^* females were crossed with Zp3-Cre^tg/+^ males. *Prkci^+/-^ ;Prkcz^-/-^* animals were generated by mating *Prkci^flox/flox^ ; Prkcz ^-/-^* females with *Prkcz*^-/-^ ;Zp3-Cre^tg/+^ males. *Rac1^flox/flox^*females were crossed with Zp3-Cre^tg/+^ males to generate *Rac1^+/−^*animals. Standard tail genotyping procedures were used to genotype transgenic mice (for primers and PCR product sizes, see Table S1). *Prkci^+/-^ ;Prkcz^-/-^* embryos were generated by crossing *Prkci ^+/-^ ;Prkcz ^-/-^* females with *Prkcz^-/-^* males. *Rac1^+/−^, Rac1^-/-^,* and WT embryos were obtained by mating *Rac1^+/−^*females with *Rac1^+/−^* males. *Myh9^+/-^* embryos were obtained by mating *Myh9^+/-^* females with WT males.

### Single embryo genotyping

Transgenic mutant embryos were genotyped retrospectively after imaging. Single embryos were transferred using a mouth-pipette from the imaging dish into PCR tubes containing 10μl lysis buffer composed of PCR buffer (Fermentas, EP0402) supplemented with 0.2 mg/ml Proteinase K (Sigma, P8811). Embryos in the lysis buffer were incubated at 55°C for 1 hour and then at 96°C for 10 minutes. 3-4μl of the resulting lysate containing genomic DNA was mixed with the relevant primers (Table S1) for determination of genotype via PCR.

PCR products were mixed with 6x loading dye (Life Technologies, R0611) and were separated by electrophoresis in 1-1.2% (w/v) agarose gel (Lonza 50004) supplemented with 0.03μl/ml DNA staining dye (Serva, 39804.01) in TAE buffer. DNA fragments were visualised under ultraviolet light on a video-based gel documentation system (Intas, GEL Stick “Touch”), and fragment lengths were measured against a standardised DNA ladder (Life Technologies, SM0323, SM0313).

**Table S1:**
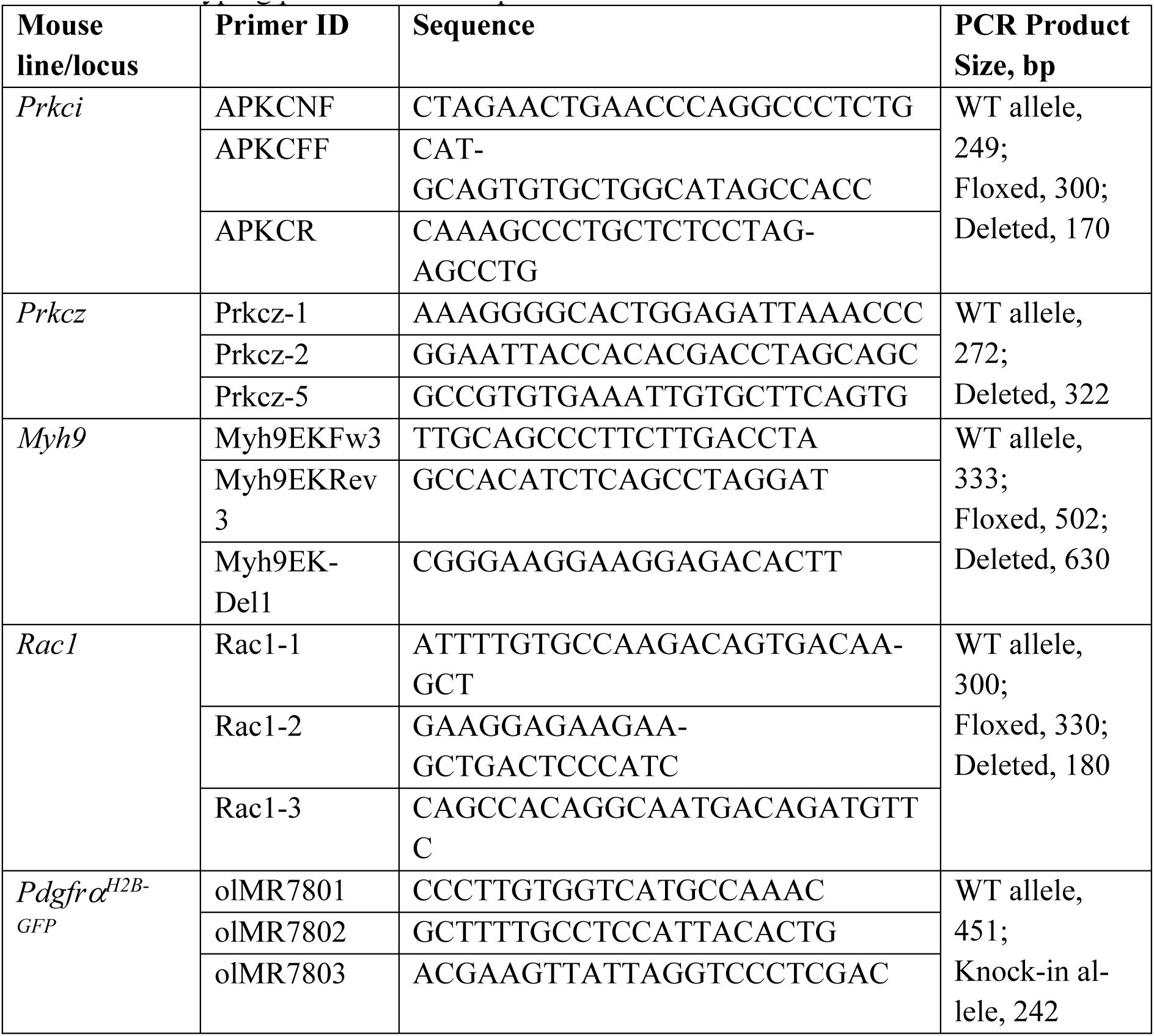
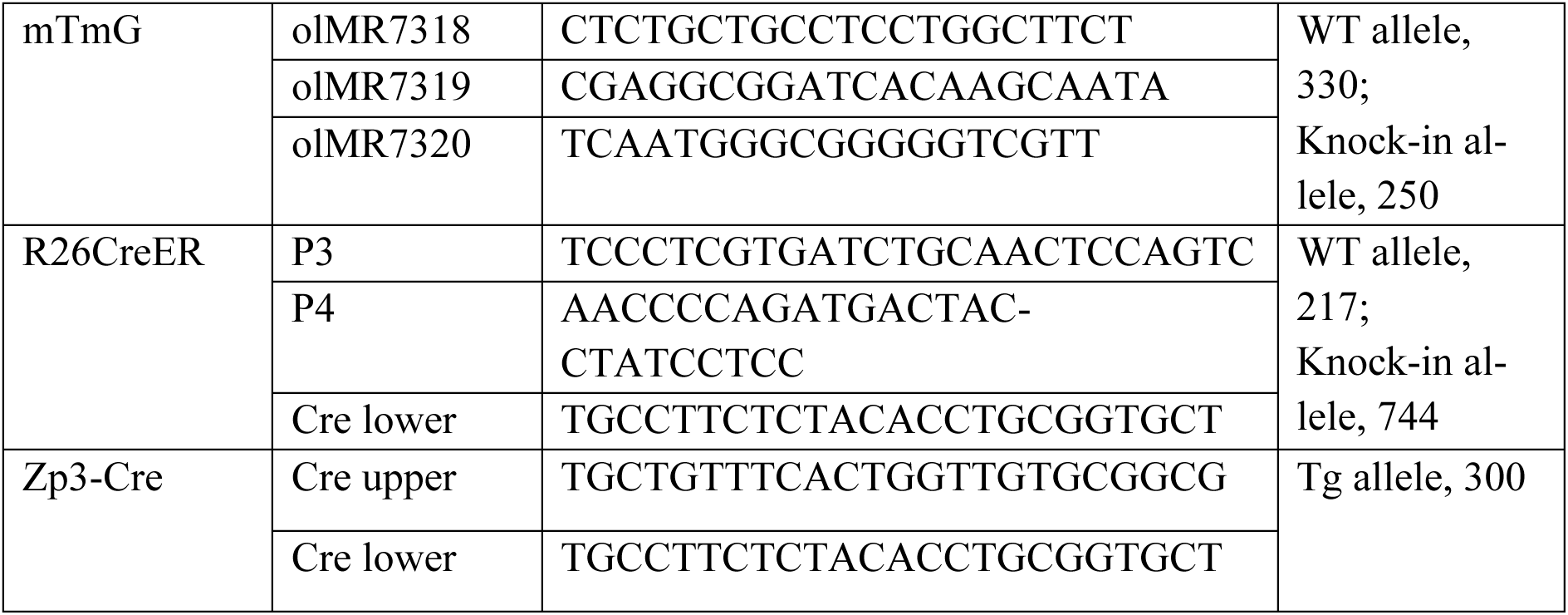
Genotyping primers and PCR product sizes.

### Intracytoplasmic sperm injection (ICSI) into monkey oocytes

Monkey oocyte collection was performed as described previously (Tsukiyama et al. 2019). Briefly, two weeks after the subcutaneous injection of 0.9 mg of a gonadotropin-releasing hormone antagonist (Leuplin for Injection Kit; Takeda Chemical Industries), a micro-infusion pump (iPRECIO SMP-200, ALZET Osmotic Pumps) with 15 IU/kg human follicle-stimulating hormone (hFSH, Gonal-f; Merck Biopharma) was embedded subcutaneously under anaesthesia and injected 7 µL/h for 10 days. After the hFSH treatment, 400 IU/kg human chorionic gonadotropin (hCG, Gonatropin; Asuka Pharmaceutical) was injected intramuscularly. Forty hours after the hCG treatment, oocytes were collected by follicular aspiration using a laparoscope (Machida Endoscope, LA-6500). Cumulus-oocyte complexes (COCs) were recovered in alpha modification of Eagle’s medium (MP Biomedicals, Solon), containing 10% serum substitute supplement (Irvine Scientific). The COCs were stripped off cumulus cells with 0.5 mg/ml hyaluronidase (Sigma Chemical). ICSI was carried out on metaphase II (MII)-stage oocytes in mTALP containing HEPES with a micromanipulator. Fresh sperm were collected by electric stimulation of the penis with no anaesthesia. Following ICSI, embryos were cultured in CMRL 1066 Medium (Thermo Fisher Scientific) supplemented with 20% FBS at 38°C in 5% CO_2_ and 5% O_2_.

### Embryo recovery and *in vitro* culture

To obtain pre-implantation embryos at different stages of development, female mice were superovulated by intraperitoneal injection of 7.5 international units (IU) of pregnant mare’s serum gonadotropin (PMSG, Intervet, Intergonan) followed by 7.5 IU of human chorionic gonadotropin (hCG; Intervet, Ovogest 1500) 48 hours later. Superovulated females were mated with male mice directly after hCG injection.

Embryos were collected from superovulated females either 68 hours post-hCG injection for uncompacted 8-cell stage, or 96 hours post-hCG injection for the 64-cell blastocyst stage, considered as the E3.5 stage. Recovery of embryos was performed under a stereomicroscope (Zeiss, StreREO Discovery.V8) equipped with a thermo plate (Tokai Hit) heated to 37°C. Oviducts and uterine horns were dissected out of female mice and submerged in global® embryo culture medium containing HEPES (LifeGlobal, LGGH-050) in 1.5ml Eppendorf tubes. Dissected oviducts and uteri were laid on a 35mm petri dish (Falcon, 351008) under the stereomicroscope and embryos were flushed out using a flushing needle attached to a 1ml syringe filled with global® medium with HEPES. After flushing, embryos were washed, transferred to 10μl drops of global® medium (LifeGlobal, LGGG-050) covered with mineral oil (Sigma, M8410) on a tissue culture dish (Falcon, 351001) and then cultured in a CO_2_ incubator (Thermo Scientific, Heracell 240i) at 37°C with 5% CO_2_.

### Immunosurgery

Zona pellucida (ZP) was removed from blastocysts with pronase (0.5% w/v Proteinase K, Sigma P8811, in global® medium containing HEPES supplemented with 0.5% PVP-40, Sigma, P0930) treatment for 2-3 minutes at 37°C. Blastocysts were washed in 10μl droplets of global® medium (LifeGlobal, LGGG-050). To isolate the inner cell mass, blastocysts were incubated in serum containing anti-mouse antibody (Cedarlane, CL2301, Lot no. 049M4847V) diluted 1:3 with global® medium for 30 minutes at 37°C. Following 2-3 brief washes in global® medium with HEPES, embryos were incubated in guinea pig complement (Sigma, 1639, Lot no. SLBX9353) diluted with global® medium in a 1:3 ratio for 30 minutes at 37°C. Lysed outer cells and remaining debris were removed by gentle pipetting with a narrow glass capillary (Brand, 708744) to isolate the inner cell mass. The isolated ICMs were cultured in 10μl drops of global® medium in a petri dish (Falcon, 351008) covered with mineral oil (Sigma, M8410) and incubated at 37°C with 5% CO_2_ for up to 24 hours.

### Embryo size manipulation

Embryos were recovered at the 8-cell stage and the zona pellucida was removed by incubation in pronase at 37°C for 2-3 minutes. For generating small-sized embryos, uncompacted 8-cell stage morulae were dissociated into the desired fraction of blastomeres by incubation in KSOM medium without Ca^2+^ and Mg^2+^ (Korotkevich et al. 2017) for 5 minutes at 37°C, followed by gentle pipetting through a narrow, flame-polished glass capillary (Brand, 708744). For generating large-sized embryos, the desired number of embryos were aggregated at the uncompacted 8-cell stage and cultured together in a single microdroplet of global® medium under mineral oil (Sigma, M8410). Embryo aggregation was encouraged by placing them in contact with each other in micro-indented wells in 3.5mm petri dishes (Falcon, 351008), ensuring that the embryos adhered to each other without drifting apart. Size-manipulated embryos were cultured until day E4.5, when a clear blastocyst cavity and coherent ICM were discernible. Embryos that failed to aggregate or showed the presence of multiple fluid cavities were discarded from further analysis.

### Generation of large embryos with mosaic labelled cells

Wild-type and fluorescent embryos expressing reporters *Pdgfra^H2B-GFP^*; mTmG were recovered at the uncompacted 8-cell stage and used for making chimeras. Fluorescent chimeras were made by pre-screening embryos for presence of the *Pdgfra^H2B-GFP^* and mTmG reporters under an inverted Zeiss Observer Z1 microscope. Embryos expressing reporters were chosen and each fluorescent embryo was mixed with 2 wild-type embryos in micro-indented wells in 3.5mm petri dishes (Falcon, 351008). The micro-wells were made in 10μl global® medium droplets covered with mineral oil (Sigma, M8410) for 24 hour *in vitro* culture until E3.5 blastocyst stage. Embryos that formed successful aggregates and showed a singular, expanded blastocyst cavity were chosen and screened for mosaic labelling of cells, and used for further live-imaging and analysis.

### Generation of mosaic labelled ICMs

Embryos were recovered from a cross between mTmG and R26CreER mouse lines at the uncompacted 8-cell stage and subject to zona pellucida removal using pronase. Embryos were then incubated in 10μM 4-hydroxytamoxifen in global® medium for 10 minutes at 37°C for tamoxifen-induced Cre-loxP recombination. The embryos were washed 5-6 times in global® medium and cultured for 24 hours in microdroplets of global® medium covered with mineral oil. At E3.5 stage, the embryos were screened for sparse conversion of mT to mG under an inverted Zeiss Observer Z1 microscope and selected for further experimental procedures. Immunosurgery was performed at E3.5 stage on the selected embryos as described earlier and the isolated ICMs were used for live-imaging.

### Micropipette aspiration

Micropipette aspiration was performed as described previously (Maître et al. 2015) to measure surface tension of ICM cells. In brief, a microforged micropipette coupled to a microfluidic pump (Fluigent, Microfluidic Flow Control System) was used to measure the surface tension of ICM cells. Micropipettes were prepared from glass capillaries (Warner Instruments, GC100T-15) using a micropipette puller (Sutter Instrument, P-1000) and a microforge (Narishige, MF-900). A fire-polished micropipette with diameter ∼7-8μm was mounted on an inverted Zeiss Observer Z1 microscope with a CSU-X1M 5000 spinning disc unit, and its movement was controlled by micromanipulators (Narishige, MON202-D). Samples were maintained at 37°C with 5% CO_2_. A stepwise increasing pressure was applied on ICM surface cells using the microfluidic pump and Dikeria software (LabVIEW), until a deformation with the same radius as that of the micropipette (R_p_) was reached. The equilibrium aspiration pressure (P_c_) was measured, images were acquired in this configuration and then the pressure was released. Care was taken to avoid aspirating the cell nuclei, since aspiration of stiffer cell nuclei may introduce artefacts in measurements. At steady state, the surface tension γ of the cells is calculated based on Young–Laplace’s law: γ = P_c_/2(1/R_p_ - 1/R_c_), in which P_c_ is the net pressure used to deform the cell of radius R_c_. Image analysis and measurement of the pipette radius R_p_ and R_c_ was done in FIJI, and calculation of surface tension was done using custom scripts in Python v3.9.

### Pharmacological inhibition

LatrunculinB (Sigma, 428020) was reconstituted in DMSO at a stock concentration of 100mM, and a final concentration of 1μM was used. CK-666 (Sigma, 182515) was resuspended in DMSO at 25mM, and a working concentration of 2μM was used. NSC23766 (Sigma, SML0952) was resuspended in DMSO at a stock concentration of 10mM, and working concentrations of 50μM, 100μM and 200μM were used. Gö6983 (Sigma, 365251) was resuspended in DMSO at 10mM and a final concentration of 5μM was used. For working concentrations of the inhibitors, respective stock concentrations were diluted in global® medium. Embryos or isolated ICMs were incubated with the appropriate working concentrations of LatrunculinB, CK-666, Gö6983, or NSC23766 and corresponding controls in μ-Slide chambered coverslips (Ibidi, 81506) before fixation in 4% PFA (refer to Immunofluorescence staining).

### Immunofluorescence staining

Mouse embryos or isolated ICMs were fixed in 4% PFA (Sigma, P6148) at room temperature for 15 minutes. Fixed embryos were washed 3 times (5 minutes each) in wash buffer DPBS Tween containing 2% BSA (Sigma, A3311), and permeabilised at room temperature for 20 minutes in permeabilisation buffer 0.5% Triton-X in DPBS; (Sigma T8787). After permeabilisation, samples were washed (3 x 5 minutes), followed by incubation in blocking buffer DPBS Tween20 (Sigma, P7949) containing 5% BSA, either overnight at 4°C or for 2 hours at room temperature. Blocked samples were then incubated with desired primary antibodies overnight at 4°C, washed (3 x 5 minutes), and incubated in fluorophore-conjugated secondary antibodies and dyes at room temperature for 2 hours. Stained samples were washed (3 x 5 minutes) and incubated in DAPI solution (Invitrogen, D3571; diluted 1:1000 in DPBS) for 10 minutes at room temperature. These samples were then transferred into individual droplets of DPBS covered with mineral oil on a 35mm glass bottom dish (MatTek, P35G-1.5-20-C) for imaging. Primary antibodies against GATA6 (R&D systems, AF1700), GATA4 (R&D systems, BAF2606), SOX2 (Cell Signaling, 23064), bi-phosphorylated myosin regulatory light chain (ppMRLC) (Cell Signaling, 3674), and Laminin (Novus Biologicals, NB300-14422) were diluted at 1:200. Primary antibodies against NANOG (ReproCell, RCAB002P-F), PKCλ (Santa Cruz Biotechnology, sc-17837), PKCζ (Santa Cruz Biotechnology, sc-17781), Integrinβ1 (Millipore, MAB1997), and active Integrinβ1 (9EG7, BD Bioscience, 553715) were diluted at 1:100. Secondary antibodies donkey anti-goat IgG Alexa Fluor 488 (Invitrogen, A11055), donkey anti-rabbit IgG Alexa Fluor 546 (Invitrogen, A10040),donkey anti-mouse IgG Alexa Fluor 555 (Invitrogen, A31570, donkey anti-rabbit IgG Alexa Fluor 647 (Invitrogen, A31573, donkey anti-mouse Cy5 (Jackson ImmunoResearch, 715-175-150), donkey anti-rat Cy5 (Jackson ImmunoResearch, 712-175-153) were used at 1:200. Immunofluorescence samples were imaged on a Zeiss LSM880 microscope with AiryScan Fast mode. A 40× water-immersion Zeiss C-Apochromat 1.2 NA objective was used, and raw Airyscan images were acquired and processed using the ZEN black software (Zeiss).

Monkey embryos that successfully developed to blastocysts were fixed between day 7-8 post ICSI in 4% paraformaldehyde (PFA, Wako 166-23251) in DPBS for 15 min at room temperature, permeabilized in DPBS with 0.5% TritonX-100 (Nacalai, 12967-32) for 30 minutes at room temperature, and blocked overnight at 4°C in DPBS with 3% BSA (Sigma, A9647) and 0.05% TritonX-100. Embryos were then transferred into primary antibody solution in blocking buffer and incubated overnight at 4°C. Primary antibodies against Alexa Fluor 647 conjugated Oct3/4 (Santa Cruz, sc-5279 AF647) and Gata4 (Cell Signalling, 36966S) were diluted at 1:200. Embryos were then washed four times for 5 minutes in blocking buffer and transferred into secondary antibody solution in blocking buffer for 2 hours at room temperature. Secondary antibody conjugated with Alexa Fluor Plus 488 against rabbit IgG (Thermo Fisher Scientific, A32790) was diluted at 1:200. Alexa Fluor Plus 555 Phalloidin (Thermo Fisher Scientific, A30106) and DAPI (Thermo Fisher Scientific, D3571) were added in the secondary antibody solution diluted at 1:400. Embryos were mounted in 1 µL drops of DPBS for imaging. Imaging of immunostained monkey embryos was performed with LSM 980 (Zeiss) with Airyscan 2 Multiplex CO-8Y mode. LD LCI Plan-Apochromat 25x/0.8 water immersion objective (Zeiss) was used.

### Time-lapse imaging

Embryos or ICMs were placed into global® medium drops covered with mineral oil on a glass bottom imaging dish (MatTek, P50G-1.5-14-F). For drug treatment experiments, embryos were placed in 60 μl of global® medium supplemented with inhibitor in 15-well glass-bottom dishes (Ibidi, 81501). Time-lapse imaging of live, fluorescent samples was performed on an inverted Zeiss Observer Z1 microscope with a CSU-X1M 5000 spinning disc unit. Excitation was achieved using 488 nm, and 561 nm laser lines through a 63/1.2 C Apo W DIC III water immersion objective. Emission was collected through 525/50 nm, 605/40 nm, band pass filters onto an EMCCD Evolve 512 camera. Images were acquired every 20 minutes for up to 12 hours. The microscope was equipped with a humidified incubation chamber to keep the sample at 37°C and supply the atmosphere with 5% CO_2_.

### Confocal live-imaging

For counting of cell numbers before and after immunosurgery, E3.5 embryos after zona removal were incubated in global® medium with 5μg/mL Hoechst 33342 (Invitrogen, H21492) for 10 minutes at 37°C, washed, and mounted in global® medium drops covered with mineral oil on 35mm glass-bottom dishes (MatTek, P35G-1.5-20-C). A full confocal z-stack was obtained for each blastocyst on the LSM 880 confocal microscope, with samples maintained in the humidified incubation chamber at 37°C and 5% CO_2_. Images were acquired with Airyscan Fast mode to reduce phototoxicity. Next, immunosurgery was performed on these blastocysts keeping track of the serial number of the embryo. Finally, confocal z-stacks of the isolated ICMs were acquired after immunosurgery with identical imaging conditions as the blastocysts.

Large-sized blastocysts with mosaic-labelled cells were transferred to individual 10μl global® medium drops covered with mineral oil on 35mm glass-bottom dishes (MatTek, P35G-1.5-20-C). The imaging dish was mounted on a Zeiss LSM 880 microscope. Embryos were maintained in a humidified chamber at 37°C and atmosphere was supplemented with 5% CO_2_. Confocal z-stacks were obtained at 20 minute intervals for up to 24 hours.

### Nuclear detection and tracking in isolated ICMs

For ICMs isolated from mouse blastocysts, nuclear detection and tracking of cell centres was performed with a semi-automatic analysis pipeline developed by (Fabrèges et al. 2023). In brief, centres of all nuclei were detected from time-lapse images using a Difference of Gaussians algorithm (Faure et al. 2016) (DoG) using the nuclear fluorescence signal. Using the high performance computing cluster at EMBL, the best parameters for the DoG algorithm were found by a grid-search procedure which explored thousands of different configuration parameters simultaneously. The best output was manually curated and used for cell tracking with a nearest neighbour algorithm (Fabrèges et al. 2023). Manual curation from the E3.5 to E4.0 stage of the ICM was performed and validated using the software Mov-IT by one operator. Cell fate was assigned based on the *Pdgfr⍺^H2B-GFP^* fluorescence intensity. All the cells in the ICM were inspected, cells that could not be traced with confidence (<1% cells) were excluded from the lineage trees.

### Evaluation of cell dynamics in isolated ICMs

Directionality analysis of cell movements from the 4D live-imaging datasets of isolated ICMs was done using cell tracking information. Cell positions in 3D were obtained for each timepoint using the Difference of Gaussians algorithm and nearest neighbour algorithm as mentioned previously. 3D cell tracking was converted into 1D radial cell positions. First, the geometric centroid of the ICM was calculated for each timepoint as the average of the x, y, z coordinates of the ICM cells. Next, radial distances of cells were calculated from the centroid at each timepoint using the Euclidean distance formula. Cell displacements were calculated for both EPI and PrE cells as the difference in radial position between two consecutive time-points. Positive displacements along the radial axis were considered as outward movement, and negative displacements as inward movement. Displacements were binned according to radial cell position and time.

### Simulations of Poissonian cellular Potts models

A cellular Potts model of the ICM system was constructed over a grid of voxels with resolution 1μm^3^. As described in a companion paper (Belousov et al. 2023, *in preparation*), the time evolution of the system was implemented as a Poissonian process, which explicitly introduces the physical time into cellular-Potts simulations through state-transition rates determined by the cellular-Potts Hamiltonian and novel kinetic parameters controlling the diffusive mobility of cells. Computer simulations were carried out with a discrete time step of 0.1 minutes. To prevent cell fragmentation, we adopted the approach described in (Durand and Guesnet 2016). The Hamiltonian of our cellular Potts model reads *E* = Σ*_ij_* J_ij_ / 2+ (κ / 2) Σ*_c_* (*V_c_* – *V_c_*)^2^, in which the first term sums over all pairs of voxels *i* and *j*, and the second term runs over individual cells *c.* The symmetric coefficients J*_ij_*=J*_ji_* vanish when the voxels *i* and *j* are not within each other’s Moore neighbourhood (Durand and Guesnet 2016), and assume one of the values listed in Table S2 depending on the voxels’ types. As described in Belousov et al. 2023 *(in preparation)*, the constants J*_medium:EPI_* and J_medium:PrE_ are chosen to correspond to the maximum and minimum surface tensions observed experimentally between medium and EPI cells, and between medium and PrE cells, respectively. Smaller surface tension differences led to lower sorting scores than those observed in the experiments, further supporting the presence of additional sorting mechanisms, as we discuss in the main text.

The total area of cells is not constrained in our simulations, in agreement with the actomyosin cortex being the main determinant of cellular shape on the relevant timescales (Graner and Riveline 2017; Salbreux, Charras, and Paluch 2012). The Poissonian transition rates are determined by kinetic parameters of *action rates* α, which control frequency of updates for one of the three voxel types (medium, EPI, and PrE) as described in Belousov et al. 2023, (*in preparation)*.

The cells’ preferred volume grows linearly as *V_c_*(t) = *V_c_*(0) + g *t* with a constant rate g until a target division volume is reached. The cell is then divided into two daughter cells along the plane perpendicular to the longest axis of the cell’s gyration tensor. The target division volume is sampled from an empirical distribution constructed from experimental volumes of mitotic cells: 2406.63, 2428.09, 2455.23, 2517.92, 2994.28, 3002.65, 3110.31, 3116.73, 3294.93, 4133.67 μm^3^. Table S2 summarizes all the numerical parameters of our simulations. For further details, see Belousov et al. 2023, *(in preparation)*.

**Table S2.** Parameter values of Poissonian cellular Potts model (Belousov et al. 2023, *in preparation*).

| Parameter | Value |
| --- | --- |
| $J_{\text{medium:EPI}}$ | $30.0 \times 10^3 k_B T$ |
| $J_{\text{medium:PrE}}$ | $9.2 \times 10^3 k_B T$ |
| $J_{\text{EPI:EPI}}$ | $18.5 \times 10^3 k_B T$ |
| $J_{\text{EPI:PrE}}$ | $25.4 \times 10^3 k_B T$ |
| $J_{\text{PrE:PrE}}$ | $13.8 \times 10^3 k_B T$ |
| $\kappa$ | $1154 k_B T / \mu\text{m}^6$ |
| $g$ | $3.36 \mu\text{m}^3 / \text{min}$ |
| $\alpha_{\text{medium}}$ | $0.95 \text{ min}^{-1}$ |
| $\alpha_{\text{EPI}}$ | $0.69 \text{ min}^{-1}$ |
| $\alpha_{\text{PrE}}$ | $2.12 \text{ min}^{-1}$ |

## Image Analysis

### Cell counting

Cell counting from immunofluorescence images of both blastocysts and isolated ICMs was done using Imaris v9.7.2 (Bitplane). The Spots module was used to detect all nuclei based on DAPI signal. Estimated nucleus diameter was set to 7μm, the automated spot detection algorithm was used to detect all cells, followed by manual validation. Cells were classified as either EPI or PrE based on the fluorescence intensities of transcription factors NANOG and GATA6 until stage E4.0, and transcription factors SOX2 and GATA4 beyond stage E4.0, respectively.

### Estimation of sorting scores

For live-imaging datasets, EPI and PrE cell positions were obtained in 3D from the nuclear detection and tracking as described previously. For immunofluorescence images, cell positions of EPI and PrE were obtained in 3D after nuclear detection using the Spots module on immunofluorescence images in Imaris. The geometric centroid of isolated ICMs was calculated as an average of the *x*, *y* and *z* coordinates of all cells in the ICM. The radial distance of ICM cells from the centroid was calculated as the Euclidean distance between the 3D coordinates of cells and the ICM centroid. Next, the sorting score was calculated as the average of the sign of the pair-wise difference between EPI and PrE radial distances.

### Unbiased classification of cell fates

Unbiased classification of cell fates into EPI and PrE from immunofluorescence images of isolated ICMs between stages E3.5 - E4.0 was done using k-means clustering in Python v3.9 with the scikit-learn library. In brief, imaging datasets were generated at the specified timepoints using identical immunostaining and imaging conditions. Fluorescence intensities of DAPI and transcription factors GATA6 and NANOG were obtained after Spot detection in Imaris. To compensate for fluorescence intensity decay with image depth, fluorescence intensities were log-transformed and a linear regression was fitted to the log values as a function of z-depth. Next, a k-means clustering algorithm was performed on the z-corrected log intensity values of GATA6 and NANOG to classify cell fates.

### Cell tracking of mosaic-labelled cells

Cell tracking of mosaic-labelled cells was performed in FIJI using a custom macro. The cell centre was determined by fitting an ellipse to the membrane signal in the equatorial plane of the cell at each timepoint. Distance of the cell from the cavity interface was measured by dropping a normal to the cavity surface from the centre of the cell at each timepoint.

### Analysis of PrE cell protrusions

Length of PrE cell protrusions was measured in FIJI, where a line segment was drawn from the centre of the nucleus to the tip of the protrusion. Angle of the protrusion with respect to the cavity was measured in FIJI as the angle between two line segments – one from the centre of the cell to the tip of the protrusion, and the other from the centre of the cell, normal to the cavity surface.

### Quantification of ectopic cells in blastocysts

Number of ectopic cells was determined using Imaris 3D visualisation. Ectopic EPI cells in E4.5 blastocysts or ICMs were defined as SOX2(+):GATA4(-) cells located at the ICM-fluid interface. Ectopic PrE cells in E4.5 blastocysts or ICMs were defined as SOX2(-):GATA4(+) cells entirely lacking a contact-free surface.

### Evaluation of cell shape

Cell shape analysis to estimate aspect ratio and circularity of cells was performed using FIJI. With the freehand selection tool, EPI and PrE cell membranes were traced manually in the equatorial plane of the cell. The long and short axis of the selection was measured as length and width, respectively, and aspect ratio was calculated as the length:width ratio. The perimeter and area of the selection was used to calculate circularity as 4ν x Area/Perimeter^2^.

### Analysis of embryo chimeras

In chimeric ICMs made from WT and *Prkci^+/-^ ;Prkcz^-/-^* embryos, cells within the ICM were first detected using the Spots module in Imaris. Next, based on fluorescence signal, mT(+) and mT(-) negative cells were classified as WT and aPKC knock-out, respectively. ICM cells were also classified as either ‘surface’ or ‘inner’ depending on whether they were in contact with the fluid medium. The proportion of mT(+) surface cells out of all surface cells was then computed and plotted for both chimeric combinations.

In chimeric blastocysts made from WT and *Myh9^+/-^* embryos, cells within the ICM were first detected using the Spots module in Imaris. Next, based on fluorescence signal, mT(+) and mT(-) negative cells were classified as WT and *Myh9^+/-^*, respectively. Using Imaris 3D visualization, GATA4(-):mT(-) cells at the ICM-fluid interface were identified as ectopic EPI cells originating from the *Myh9^+/-^* population.

### Signal intensity measurements

ppMRLC intensity analysis was performed in FIJI. A freehand line was manually traced along the outer cell surface of E3.5 ICM cells, and mean fluorescence intensity of ppMRLC was measured along this curve in an equatorial section of the ICM. Further, the ppMRLC signal was normalised by nuclear DAPI intensity of the cell.

aPKC intensity analysis was performed using FIJI. A line segment was manually drawn for E3.5 ICM cells from the inner edge to the outer edge of the cell along the radial direction. The inner edge was defined as the region of the cell towards the ICM centroid, whereas the outer edge of the cell was defined as the region of the cell facing the ICM-fluid interface. Fluorescence intensity of aPKC was measured along this line segment, with segment width set to 10 pixels. The fluorescence intensity was binned into a fixed number of intervals to normalize cell length. Polarisation index was calculated as the ratio between mean aPKC fluorescence intensity over 1/4th distance from outer edge and mean aPKC intensity over 1/4th distance from the inner edge.

Laminin intensity analysis was performed in FIJI. For ICMs, laminin intensity was measured along the cell boundary of EPI and PrE precursors in the equatorial plane of the ICM. For blastocysts, a maximum intensity projection of the ICM was obtained and the laminin fluorescence intensity was measured along a line segment of width 10 pixels, drawn from the ICM-TE interface to the ICM-cavity interface. The line measurements were normalised by the maximum fluorescence intensity and then smoothed using a rolling average.

Colocalisation analysis of integrinβ1 and laminin was performed in FIJI using the Colocalization plugin. Cells of interest were cropped out of whole blastocyst images and the integrinβ1 and laminin signals were thresholded to remove background signal. Thresholded images were used as input for the plugin, to calculate Mander’s co-efficients and Pearson’s correlation coefficient for signal overlap.

### Estimation of ICM-fluid surface area and ICM volume

For size-manipulated mouse embryos and monkey embryos, physical dimensions of the ICM were measured in FIJI from immunofluorescence images of blastocysts. First, a half-circle or crescent-shaped cross-section of the ICM was generated in 2D by re-slicing 3D confocal images. The ICM base diameter was measured as the length of the line segment joining the extreme tips of the crescent shape, and ICM radius *R* as half of the diameter. The major height *H* of the ICM was measured as the largest perpendicular distance of the ICM-polar TE interface from the ICM diameter. The minor height *h* was measured as the largest perpendicular distance of the ICM fluid interface from the ICM diameter.

The volume of the ICM was measured as the difference between the volumes of two spherical caps, one with base radius *R* and height *H*, the other with base radius *R* and height *h*. The ICM fluid interface area was measured as the surface area of the spherical cap with base radius *R* and height *h*. The mathematical equations used for calculating volume and surface area of a spherical cap with base radius *r* and height *h* are as follows:

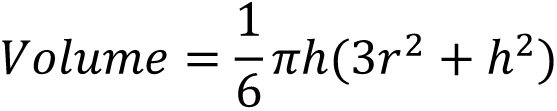

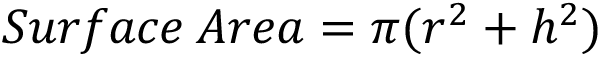

### Estimation of PrE cell apical areas

PrE cell apical area from mouse and monkey embyros was measured using immunofluorescence images of blastocysts in Napari. In brief, freehand lines were manually traced with the Labels tool along the cell-fluid interface in individual PrE cells from confocal z-stacks based on the actin signal. With the label-interpolator plugin, a surface contour was constructed for individual cells from the traced lines. Using a custom python script, the number of pixels in the surface contour were measured to estimate the cell apical area.

### Estimation of PrE monolayer, gap and multilayer probabilities

The interval of typical values for the PrE area A_PrE_ were estimated using the 10^th^ and 90^th^ quantiles of the PrE cells apical surface area A_10%_ and A_90%_, respectively. For a given total number *n* of EPI and PrE cells in the ICM, the minimal A_PrE_ was estimated as *f n* A_10%_ and maximal A_PrE_ was estimated as *f n* A_90%_, in which *f* is the fraction of PrE cells: 60% in mouse embryos, and 70% in monkey embryos. The probability of forming a PrE monolayer was determined by the relative frequency of embryos whose ICM-cavity interface area A_interface_ was within the area enclosed between minimal A_PrE_ and maximal A_PrE_. Likewise, the probability of a gap (or multilayer) was estimated as the relative frequency of embryos with A_Interface_ > maximal A_PrE_ (or A_Interface_ < minimal A_PrE_).

### Statistical Analysis

All statistical analysis and data visualization was performed in Python v3.9, with the SciPy package used for statistical testing. Normality of the distribution for datasets was tested by a Shapiro-Wilk test. If the dataset followed normal distribution, an independent-samples t-test was used for comparisons between two groups, or one-way ANOVA was used for testing more than two groups, followed by a Tukey’s post hoc test. Otherwise, a non-parametric Kruskal Wallis test or Mann-Whitney U test was used for comparisons between different groups. For testing correlation, linear regression was used and the Pearson correlation coefficient was estimated. No statistical methods were used to predetermine sample size, and no randomisation method was used. The investigators were not blinded during experiments. Box and whisker plots depict the minima, lower quartiles, medians, upper quartiles and maxima. Error bars indicate mean±SD unless mentioned otherwise. Number of embryos analysed for different experimental conditions are indicated as n-values unless mentioned otherwise. Sample sizes, statistical tests, correlation coefficients, and p-values are indicated in text, figures and figure legends.

