## Supplementary figures and legends for "Apical-driven cell sorting optimised for tissue geometry ensures robust patterning"

### SUPPLEMENTARY FIGURES, TITLES AND LEGENDS

#### Supplementary Figure 1

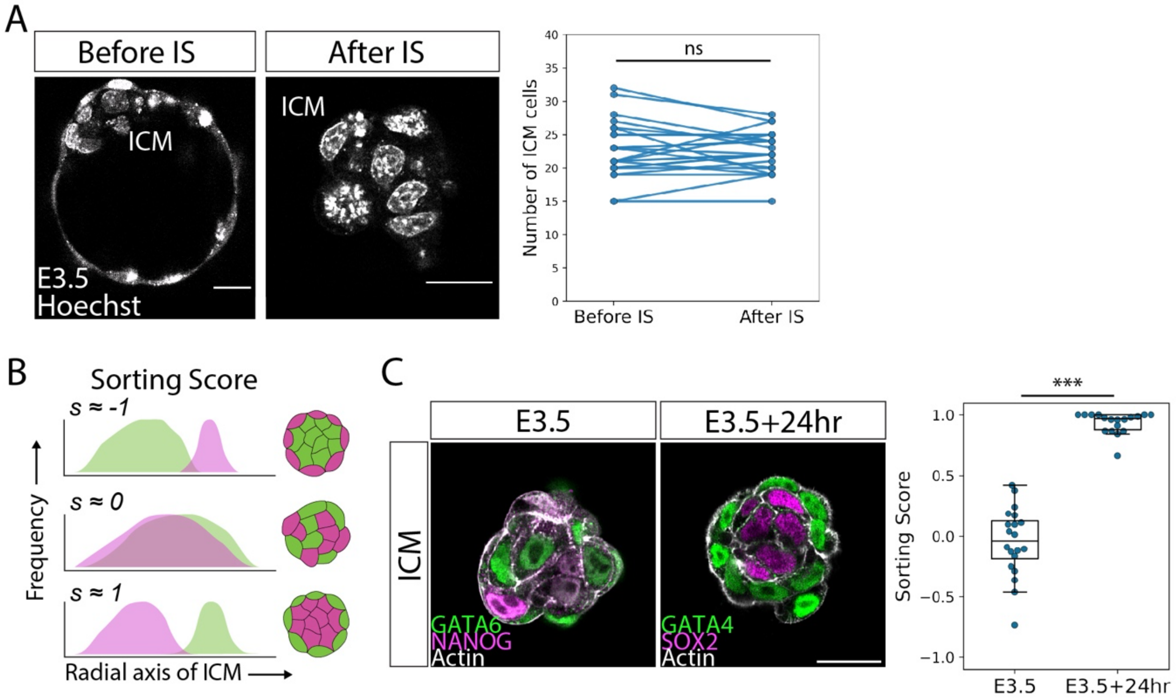

**Supplementary Figure 1: Differential cell movements between epiblast and primitive endoderm contribute to fate segregation in the ICM, related to Figure 1.**

A. Representative images of an E3.5 blastocyst and corresponding isolated ICM after immunosurgery with quantification of total cell numbers in the ICM before and after immunosurgery. Paired samples *t*-test, $p$=0.644 with n=24 embryos.
B. Schematic representation for quantification of the sorting score in isolated ICMs. Sorting score values range from s=-1 to s=1, with s=-1 indicating EPI enveloping PrE, s=0 indicating salt-and-pepper distribution of cell types, and s=1 indicating PrE enveloping EPI.
C. Representative immunofluorescence images of ICMs isolated at stage E3.5 (left), compared to ICMs isolated at stage E3.5 followed by 24-hour *in vitro* culture (right), and quantification of the sorting score in these experimental groups. Mann Whitney U-test, $p$=1.45e$^{-07}$ and n=20, 18 ICMs for the two groups, respectively.

Scale bar 20µm.

*ns, non-significant, ***$p \leq 0.001$*

**Supplementary Figure 2**

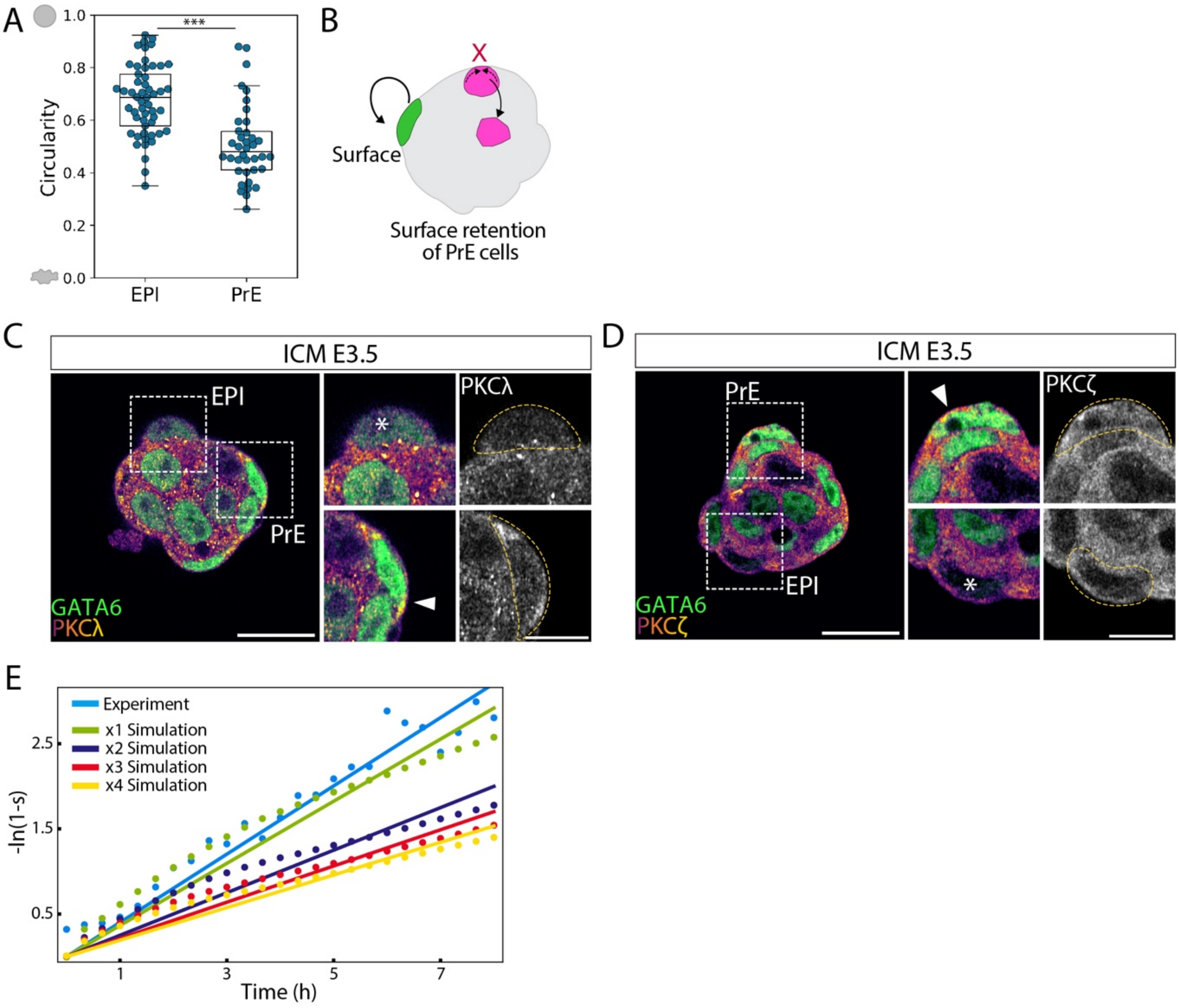

**Supplementary Figure 2: Acquisition of the apical domain decreases surface tension and is sufficient for retaining PrE cells at the fluid interface, related to Figure 2.**

A. Analysis of surface cell circularity in E3.5 isolated ICMs to compare EPI and PrE cell shape. n=53, 38 cells from 16 ICMs. Mann-Whitney U test, $p=5.23e^{-07}$
B. Schematic diagram for the retention hypothesis.
C. Representative immunofluorescence image of an isolated ICM at stage E3.5 showing distribution of PKCλ, and higher magnification images of PKCλ localisation in EPI (GATA6-low, top) and PrE (GATA6-high, bottom) cells on the ICM surface. White arrowhead, PKCλ localisation at the apical cortex in surface PrE cells. White asterisk, an EPI cell at the ICM surface. Yellow dotted line, cell boundary.
D. Representative immunofluorescence image of an isolated ICM at stage E3.5 showing distribution of PKCζ, and higher magnification images of PKCζ localisation in PrE (GATA6-high, top) and EPI (GATA6-low, bottom) cells on the ICM surface. White arrowhead, PKCζ localisation at the apical cortex in surface PrE cells. White asterisk, an EPI cell at the ICM surface. Yellow dotted line, cell boundary.
E. Relaxation time $\tau$ is obtained by fitting the exponent of the sorting-score time series $s=1-e^{-t/\tau}$, for experimental live-imaging data and simulations of isolated ICMs of different sizes. Dots indicate the data points for the time-series, solid line indicates the linear fit, and the slope of the fit is $1/\tau$.

Scale bars, 20μm; scale bars for zoomed-in panels, 10μm.

**Supplementary Figure 3**

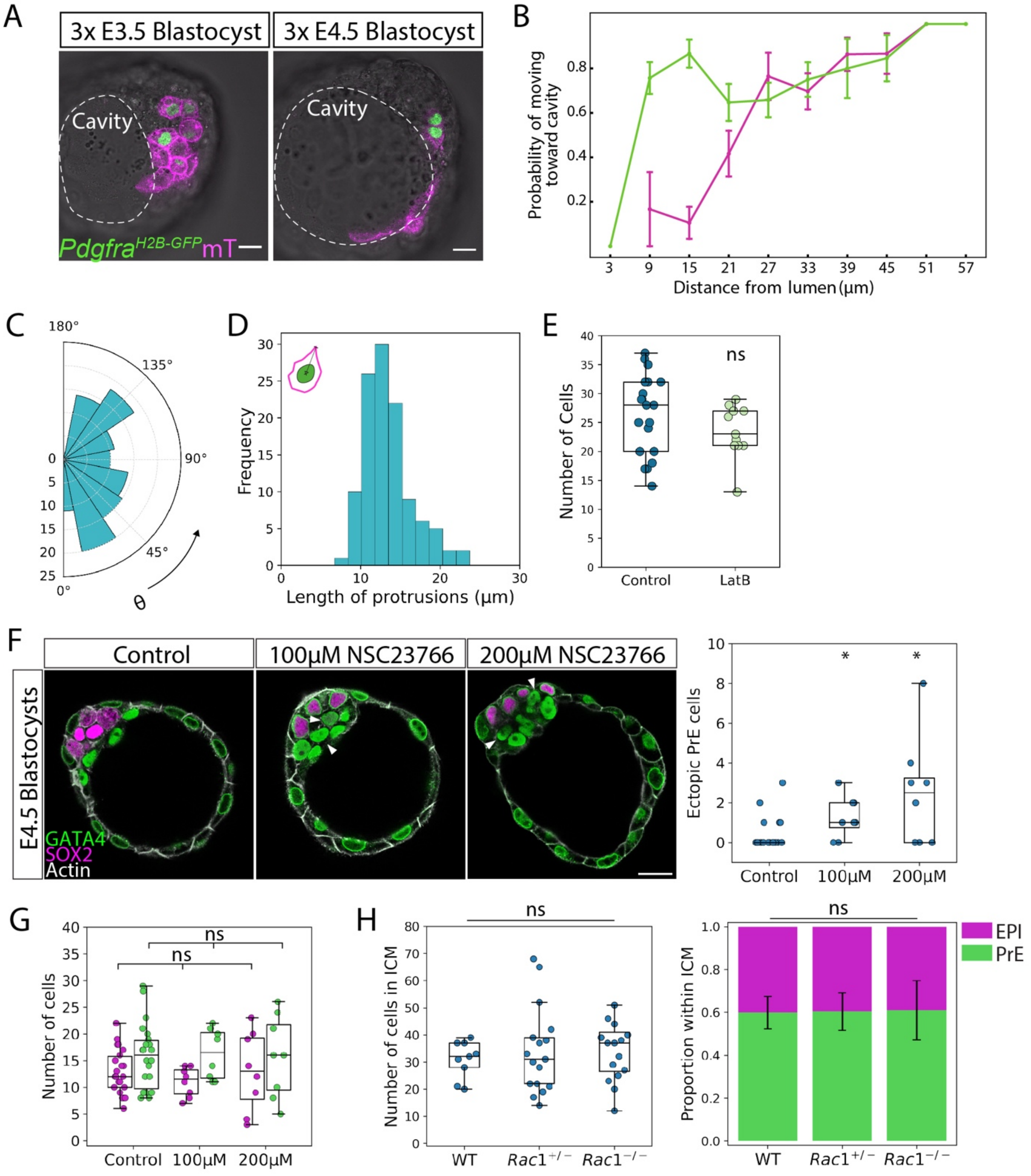

**Supplementary Figure 3: Cell sorting involves active directed migration of PrE cells towards the surface via actin-mediated protrusions, related to figure 3.**

A. Representative time-lapse images of mosaical-labelled blastocysts at E3.5 and E4.5 generated to visualise EPI and PrE cell dynamics.
B. Probability of EPI and PrE cells to move towards the cavity surface as a function of cellular distance from the lumen. Probabilities estimated from single-cell tracking in mosaic-labelled blastocysts from Figure 2B,C.
C. Polar histogram for protrusion angles sampled from a uniform distribution.
D. Distribution of protrusion length in PrE cells as measured from centre of the nucleus to tip of the longest protrusion. Mean protrusion length = 13.38 ± 3.12μm.
E. Quantification of total number of ICM cells in control and latrunculinB-treated ICMs. n=20,11 ICMs for the two groups, respectively. Mann-Whitney U test, $p = 0.207$.
F. Representative images of control E4.5 blastocysts and 100μM and 200μM NSC23766-treated E4.5 blastocysts and quantification of number of ectopic PrE cells in each condition. White arrowheads, ectopic PrE cells. n= 22, 8, 8 blastocysts for the treatment groups, respectively. Mann-Whitney U test, $p = 0.011$ for comparison between control and 100μM group, $p=0.012$ for comparison between control and 200μM group.
G. Quantification of total number of EPI and PrE cells in control and NSC23766-treated E4.5 blastocysts. n= 22, 8, 8 blastocysts respectively. Kruskal-Wallis test, $p=0.62$ for EPI cell numbers, $p=0.903$ for PrE cell numbers across the treatment groups.
H. Quantification of total number of ICM cells in WT, $Rac1^{+/-}$, and $Rac1^{-/-}$ E4.5 blastocysts. n=9, 17, 16 blastocysts respectively. Kruskal-Wallis test, $p=0.63$. Quantification of EPI/PrE cell fate proportion within the ICM in WT, $Rac1^{+/-}$, and $Rac1^{-/-}$ E4.5 blastocysts, plotted as mean ± SD. One-way ANOVA, p=0.567.

Scale bar 20μm.

*ns, non-significant, *p≤0.05*

**Supplementary Figure 4**

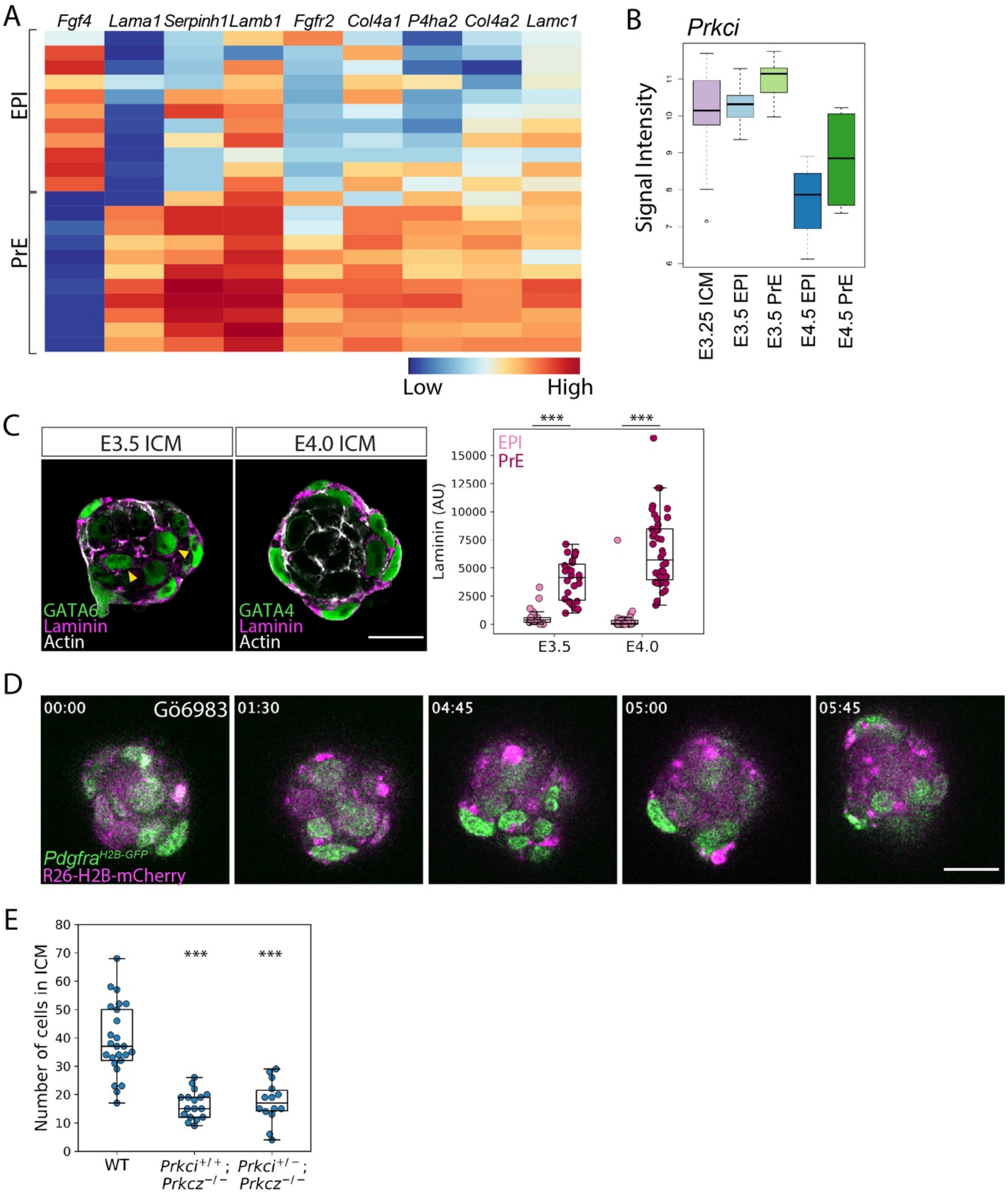

**Supplementary Figure 4: Apical polarisation in PrE cells is required for directed migration and sorting, related to Figure 4.**

A. Heatmap indicating the expression levels of extracellular matrix-related components in EPI and PrE cells from E3.5 blastocysts analysed by single-cell qPCR (22 cells from 3 embryos at E3.5).
B. Boxplot indicating differential expression of *Prkci* among EPI and PrE cells in the ICM cells at different stages.
C. Immunofluorescence images of E3.5 and E4.0 isolated ICMs stained for pan-Laminin. Quantification of laminin deposition around EPI and PrE cells in E3.5 and E4.0 isolated ICMs. n=18, 27, 37, 42 cells for the different groups, respectively. Mann-Whitney U-test, $p$=2.54e$^{-07}$ for stage E3.5, $p$=1.46e$^{-13}$ for stage E4.0.
D. Time-lapse imaging of ICMs isolated from E3.5 blastocysts expressing PrE-specific H2B-GFP ($Pdgfr\alpha^{H2B\text{-}GFP}$, green) and ubiquitous H2B-mCherry (*R26-H2B-mCherry*, magenta) and treated with pan-aPKC inhibitor Gö6983. White arrowhead marks a PrE cell unable to maintain its surface position. Time is indicated as hh:mm, t=00:00 corresponds to start of live-imaging at stage E3.5+3hours, following completion of immunosurgery.
E. Box and scatter plot for total ICM cell number in WT, $Prkci^{+/+};Prkcz^{-/-}$, and $Prkci^{+/-};Prkcz^{-/-}$ blastocysts at E4.5 stage. n=25,17 and 14 for the experimental groups, respectively. Independent samples t-test, $p$=2.23e$^{-08}$ for $Prkci^{+/+};Prkcz^{-/-}$, and $p$= 1.25e$^{-06}$ for $Prkci^{+/-};Prkcz^{-/-}$ blastocysts.

Scale bars, 20µm.

*ns, non-significant, ***p≤0.001*

**Supplementary Figure 5**

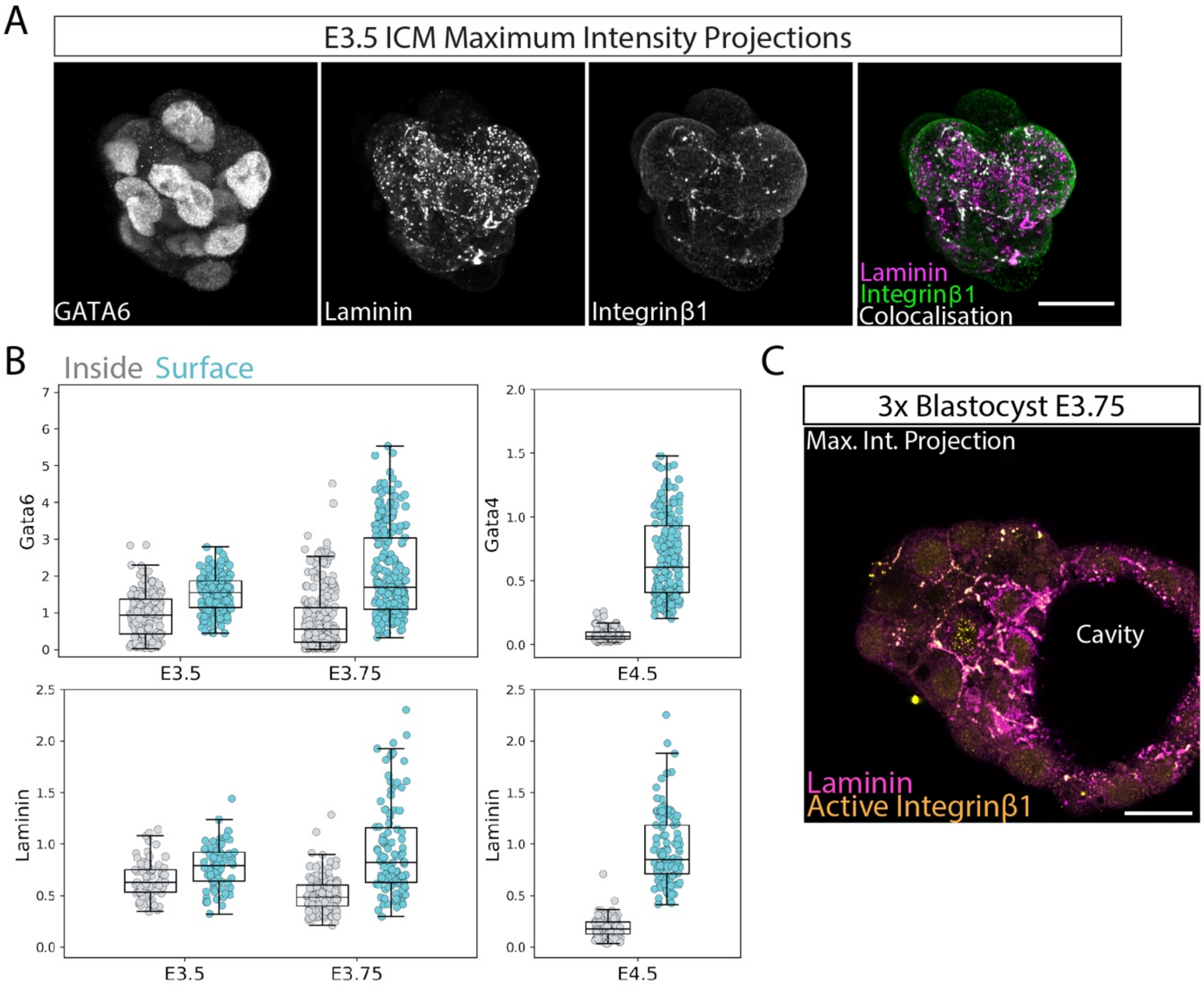

**Supplementary Figure 5: Extracellular matrix deposited in the ICM guides PrE cells towards the cavity surface, related to Figure 5.**

A. Immunofluorescence maximum intensity projection of an ICM stage E3.5 showing the distribution of laminin, integrinb1, and their colocalisation.
B. Quantification of nuclear GATA6/GATA4 fluorescence intensities (top) and laminin accumulation around cells (bottom) in the ICM cells from 3x blastocysts at stages E3.5, E3.75 and E4.5. Cells located at the ICM-cavity interface were considered 'surface' cells (cyan), whereas cells without contact with the cavity were considered 'inner' cells (grey).
C. Representative immunofluorescence image of a 3x blastocyst at stage E3.75 showing the ECM gradient towards the cavity surface in the ICM.

Scale bar 20μm.

#### Supplementary Figure 6

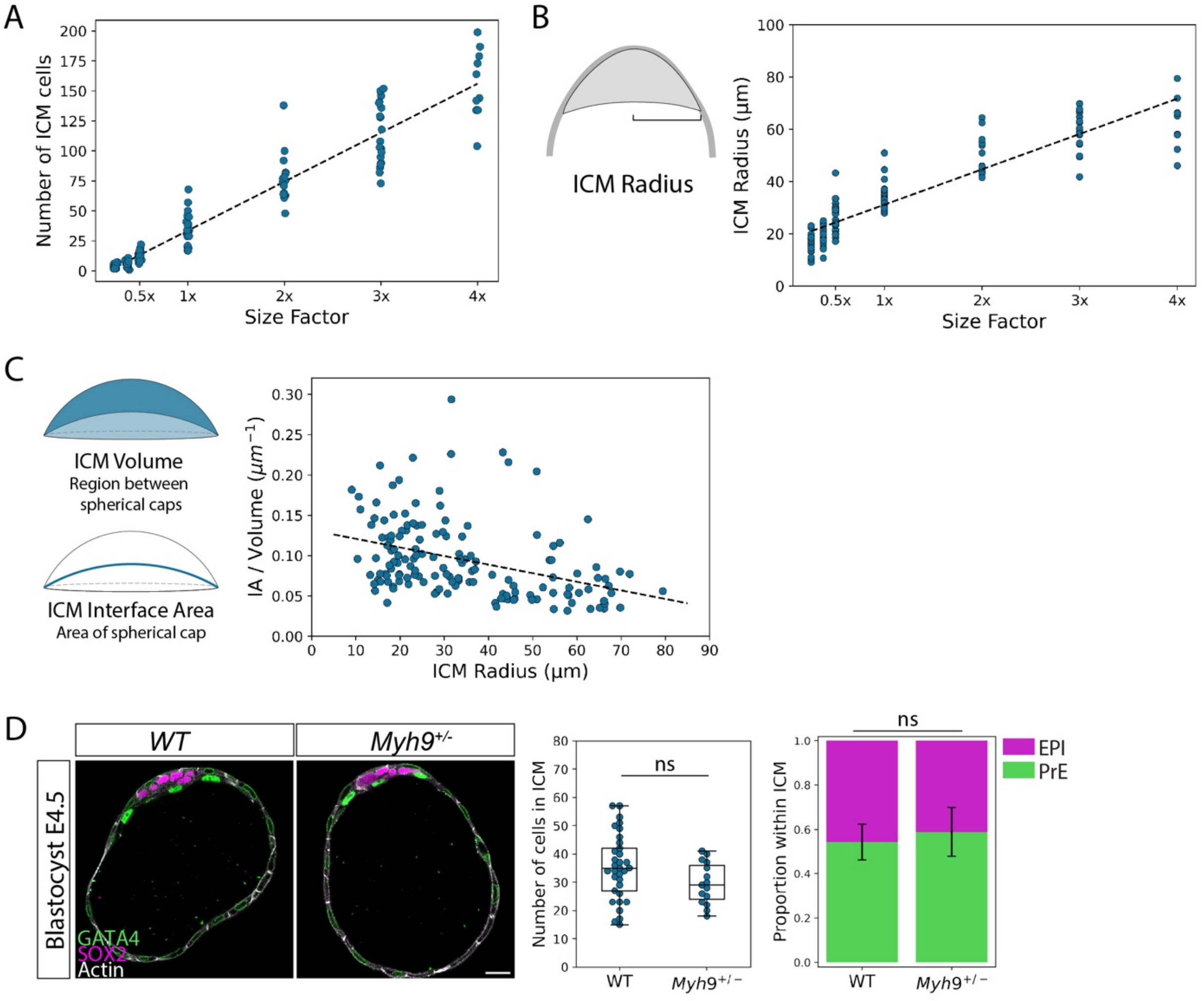

**Supplementary Figure 6: The fixed proportion of EPI/PrE cells without cell fate switching challenges precision in ICM patterning**

A. Number of cells in the ICM scales linearly with embryo size factor in size-manipulated E4.5 blastocysts. Dotted line, linear regression with Pearson's R=0.958, $p$=1.95e$^{-84}$. n = 29 embryos for 2/8x, 26 for 3/8x, 17 for 4/8x, 29 for 1x, 19 for 2x, 24 for 3x, and 10 embryos for 4x size ratios

B. ICM radius scales linearly with embryo size factor in size-manipulated E4.5 blastocysts. Dotted line, linear regression, Pearson's R=0.908, $p$=1.76e$^{-57}$.

C. ICM interface area-to-volume ratio decreases with increasing ICM radius and embryo size. Dotted line, linear regression, Pearson's R=-0.386, $p$=1.17e$^{-06}$.

D. Representative immunofluorescence images of WT and *Myh9*$^{+/-}$ blastocysts at stage E4.5. Box and scatter plot for quantification of total ICM cell number in WT and *Myh9*$^{+/-}$ blastocysts at stage E4.5. n=33, 15 embryos for WT and *Myh9*$^{+/-}$, respectively. Independent samples t-test, $p$=0.086.
Stacked bar plot indicating EPI/PrE cell fate proportion within the ICM in WT and *Myh9*$^{+/-}$ blastocysts at stage E4.5, plotted as mean ± SD. n = 33, 15 embryos for WT and *Myh9*$^{+/-}$, respectively. Independent samples t-test, $p$=0.121.

Scale bar 20µm.

*ns, non-significant*

#### Supplementary Figure 7

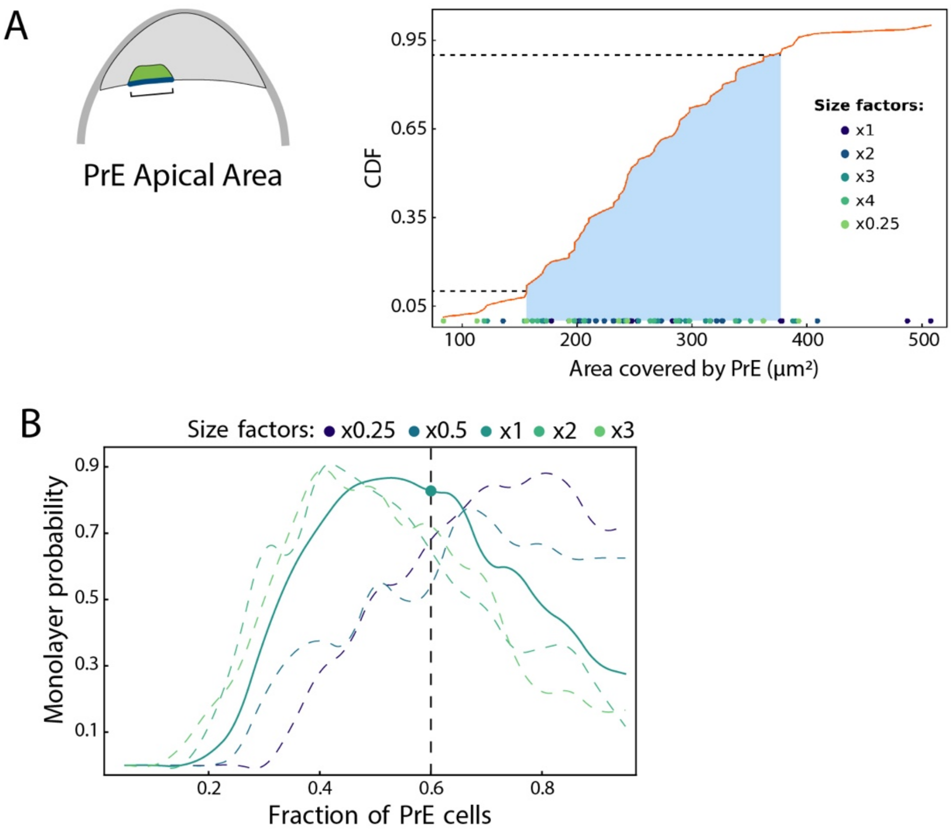

**Supplementary Figure 7: The fixed proportion of EPI/PrE cells is optimal for the specific embryo size and ICM geometry**

A. Schematic diagram for Pre cell apical area and distribution of individual PrE apical areas measured in 3D across size-manipulated E4.5 blastocysts. Blue shaded region corresponds to the region between the $10^{th}$ and $90^{th}$ percentile of apical area measurements with $q_{10\%}$=157mm$^2$ and $q_{90\%}$=376mm$^2$.

B. Probability of monolayer PrE formation for size-manipulated embryos at stage E4.5 with respect to PrE proportion within the ICM. Black dotted line indicates the fixed 60% PrE proportion in the ICM.

#### SUPPLEMENTARY VIDEOS

**Supplementary video 1**
Isolated ICMs as a reduced system to study EPI/PrE fate segregation. Time-lapse images of an isolated ICM expressing $Pdgfr\alpha^{H2B\text{-}GFP}$ (green) and *R26-H2B-mCherry* (magenta), developing *in vitro* between stages E3.5 and E4.0. Time, hours:minutes, with t=00:00 starting at stage E3.5+3hours. Scale bar, 20 µm.

**Supplementary video 2**
Mosaic-labelling of cells allows visualisation of cell shape changes in individual cells. Time-lapse images of an isolated ICM labelled in a mosaic manner via tamoxifen-induced Cre-mediated conversion of mT (magenta) to mG (green) developing *in vitro* between stages E3.5 and E4.0. Time, hours:minutes with t=00:00 starting at stage E3.5+3hours. Scale bar, 20 µm.

**Supplementary video 3**
Simulation from the Poissonian cellular Potts model. EPI/PrE sorting simulated by a Poissonian cellular Potts model, depicting the lower hemisphere of the ICM for visualisation of the transverse section. Simulation is run from stage E3.5+3hours for 8 hours, controlled at temperature 37°C. EPI is shown in magenta and PrE is shown in green.

**Supplementary video 4**
Mosaic-labelling of EPI and PrE cells allows visualisation of cell shape changes and tracking during fate segregation in the blastocyst.
Time-lapse images of mosaic-labelled 3x blastocysts with cells expressing $Pdgfr\alpha^{H2B\text{-}GFP}$ (green) and membrane td-tomato (mT, magenta) and brightfield, to visualise EPI and PrE cell dynamics. Time is indicated as hh:mm, t=00:00 corresponds to start of live-imaging at stage E3.5+3hours. Scale bar, 20 µm.

**Supplementary video 5**
Visualisation of cell shape changes in EPI cells during fate segregation in the blastocyst.
Time-lapse images of a representative EPI cell in a mosaic-labelled 3x blastocyst expressing $Pdgfr\alpha^{H2B\text{-}GFP}$ (green) and membrane td-tomato (mT, magenta) (left), with a merged bright-field view of the blastocyst (right). Cell fate is assigned as EPI due to the low $Pdgfr\alpha^{H2B\text{-}GFP}$ expression. Time is indicated as hh:mm, t=00:00 corresponds to start of live-imaging at stage E3.5+3hours. Scale bar, 20 µm.

**Supplementary video 6**
Visualisation of cell shape changes and membrane protrusions in PrE cells during fate segregation in the blastocyst.
Time-lapse images of a representative PrE cell in a mosaic-labelled 3x blastocyst expressing $Pdgfr\alpha^{H2B\text{-}GFP}$ (green) and membrane td-tomato (mT, magenta) (left), with a merged bright-field view of the blastocyst (right) to locate the ICM-fluid interface. Cell fate is assigned as PrE due to the high $Pdgfr\alpha^{H2B\text{-}GFP}$ expression. Time is indicated as hh:mm, t=00:00 corresponds to start of live-imaging at stage E3.5+3hours. Scale bar, 20 µm.

**Supplementary video 7**
Rac-1 knock-out embryos fail to segregate PrE by E4.5 stage in blastocysts. Z-stacks from immunofluorescence images of $Rac1^{+/-}$ (left) and $Rac1^{-/-}$ (right) blastocysts at stage E4.5, visualising GATA4 (green), SOX2 (magenta) and actin (grey). Scale bars, 20 µm.

**Supplementary video 8**
Inhibition of aPKC with Gö6983 results in failure of EPI/PrE segregation. Time-lapse images of an isolated ICM expressing $Pdgfr\alpha^{H2B\text{-}GFP}$ (green) and *R26-H2B-mCherry* (magenta) treated with pan-aPKC inhibitor Gö6983. Time, hours:minutes with t=00:00 starting at stage E3.5+3hours. Scale bar, 20 µm.

**Supplementary video 9**
aPKC knock-out embryos fail to segregate EPI and PrE by E4.5 stage in blastocysts. Z-stacks from immunofluorescence images of $Prkci^{+/+};Prkcz^{-/-}$ (left) and $Prkci^{+/-};Prkcz^{-/-}$ (right) blastocysts at stage E4.5, visualising GATA4 (green), SOX2 (magenta) and actin (grey). Scale bars, 20 µm.

**Supplementary video 10**
Perturbation of embryo size challenges precision of ICM patterning. Z-stacks from immunofluorescence images of a small-sized blastocyst (3/8x, left) and a large-sized blastocyst (4x, right) blastocysts at stage E4.5, visualising GATA4 (green), SOX2 (magenta) and actin (grey). Scale bars, 20 µm.

**Supplementary video 11**
$Myh9^{+/-}$ blastocysts show successfully segregate EPI/PrE cell in the ICM in E4.5 blastocysts. Immunofluorescence z-stack of an E4.5 $Myh9^{+/-}$ blastocyst with GATA4 in green, SOX2 in magenta and actin in grey. Scale bar, 20 µm.

**Supplementary video 12**
Comparison of ICM cell numbers, embryo size and ICM geometry between mouse and monkey blastocysts. Z-stacks from immunofluorescence images of an E4.5 mouse blastocyst (left) visualising GATA4 (green), SOX2 (magenta) and actin (grey), and an E7.0-E8.0 monkey blastocyst (right), visualising GATA4 (green), OCT3/4 (magenta) and actin (grey). Scale bars, 20 µm.
